# Systematic Growth Factor Profiling Platform for 3D Tumor Models Reveals Estradiol-Responsive Cellular Mechanisms of Immunotherapy Resistance

**DOI:** 10.1101/2025.09.04.674374

**Authors:** Kwanghwan Lee, Minsung Kim, Si On Lim, Dong-Ju Shin, Yun Shin, Jung-Joo Choi, Maria Lee, Hyun Ju Kang, Jeong-Won Lee, Jin-Ku Lee

## Abstract

Current organoid culture systems face critical limitations: standardized growth factor formulations fail to capture patient-specific signaling requirements, and single-cell-type approaches overlook tumor-stromal interactions essential for understanding immunotherapy resistance. To address these challenges, we developed an innovative biofabrication platform that systematically integrates patient-derived three-dimensional (3D) cultures with comprehensive growth factor profiling across 128 combinations. Through systematic screening of 23 ovarian cancer patient samples and single-cell RNA sequencing, we identified two estradiol-responsive cellular populations that coordinate immunosuppression: a malignant cell fraction (MAL.PDCD5) and a cancer-associated fibroblast (CAF) fraction (FB.TNFSF10). MAL.PDCD5 cells suppress immune infiltration by downregulating antigen presentation, while FB.TNFSF10 cells promote immunosuppression through enhanced TGF-β signaling. Spatial transcriptomic analysis revealed striking mutual exclusivity between FB.TNFSF10 cells and T/NK cells, providing evidence of active immune cell exclusion. Most significantly, FB.TNFSF10 abundance emerged as a robust predictor of immune checkpoint inhibitor therapy resistance across multiple cancer cohorts, independent of conventional biomarkers. This biofabrication platform provides a scalable framework for drug screening and biomarker discovery, with immediate applications in precision medicine for patient stratification and combination therapy development.

## 1. Introduction

The biofabrication of physiologically relevant tumor models represents a critical advancement in cancer research, enabling the recapitulation of complex cellular interactions that govern treatment responses [1, 2]. While traditional 2D cell culture systems have provided valuable insights into cancer biology, they fail to capture the three-dimensional architecture and cellular heterogeneity characteristic of native tumor microenvironments (TMEs) [3, 4]. Recent advances in biofabrication technologies, including organoid culture systems, bioprinting, and microfluidic platforms, have opened new possibilities for engineering tumor models that more accurately represent *in vivo* conditions [5–8].

However, existing 3D culture protocols face significant limitations. Most rely on empirically derived, standardized media formulations that fail to capture patient-specific growth factor requirements, using one-size-fits-all combinations without systematic optimization [9, 10]. This approach potentially misses critical signaling dependencies that vary between patients and tumor subtypes [11, 12]. Additionally, the complexity of the TME extends beyond cancer cells to include various stromal components, immune cells, and signaling molecules that collectively influence tumor progression and therapeutic responses [13, 14]. Cancer-associated fibroblasts (CAFs) have emerged as critical orchestrators of TME dynamics [15], exhibiting remarkable plasticity and adopting distinct functional states that can either promote or suppress antitumor immunity [16]. Yet current biofabrication approaches have not fully captured the regulatory mechanisms governing CAF heterogeneity, particularly in hormone-responsive contexts [17].

To address these limitations, we developed an innovative biofabrication platform that systematically integrates patient-derived organoid cultures with comprehensive growth factor profiling across 128 combinations. This approach overcomes empirical media limitations and provides unprecedented insights into patient-specific signaling requirements. We pioneered the integration of single-cell RNA sequencing with 3D culture systems to investigate estradiol-mediated immune regulation in cancer, uniquely combining patient-derived cultures, systematic growth factor screening, and molecular profiling to identify specific cellular populations that drive immunosuppression and therapy resistance. This biofabrication strategy provides a scalable framework for modeling hormone-responsive tumor microenvironments and offers new opportunities for drug screening and biomarker discovery in cancer immunotherapy.

## 2. Materials and methods

### 2.1. Reagents and tools table

#### Basal media composition and growth factors

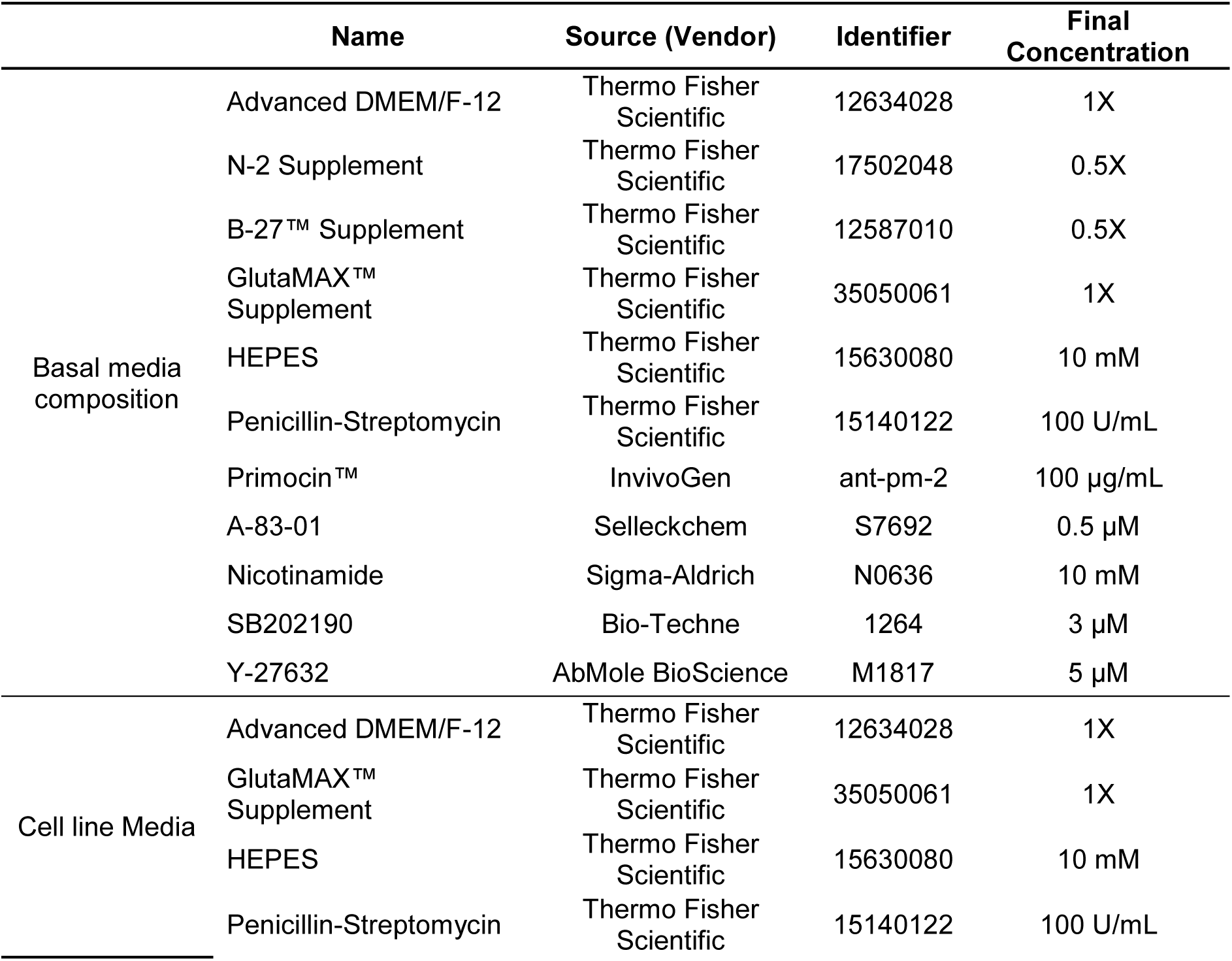
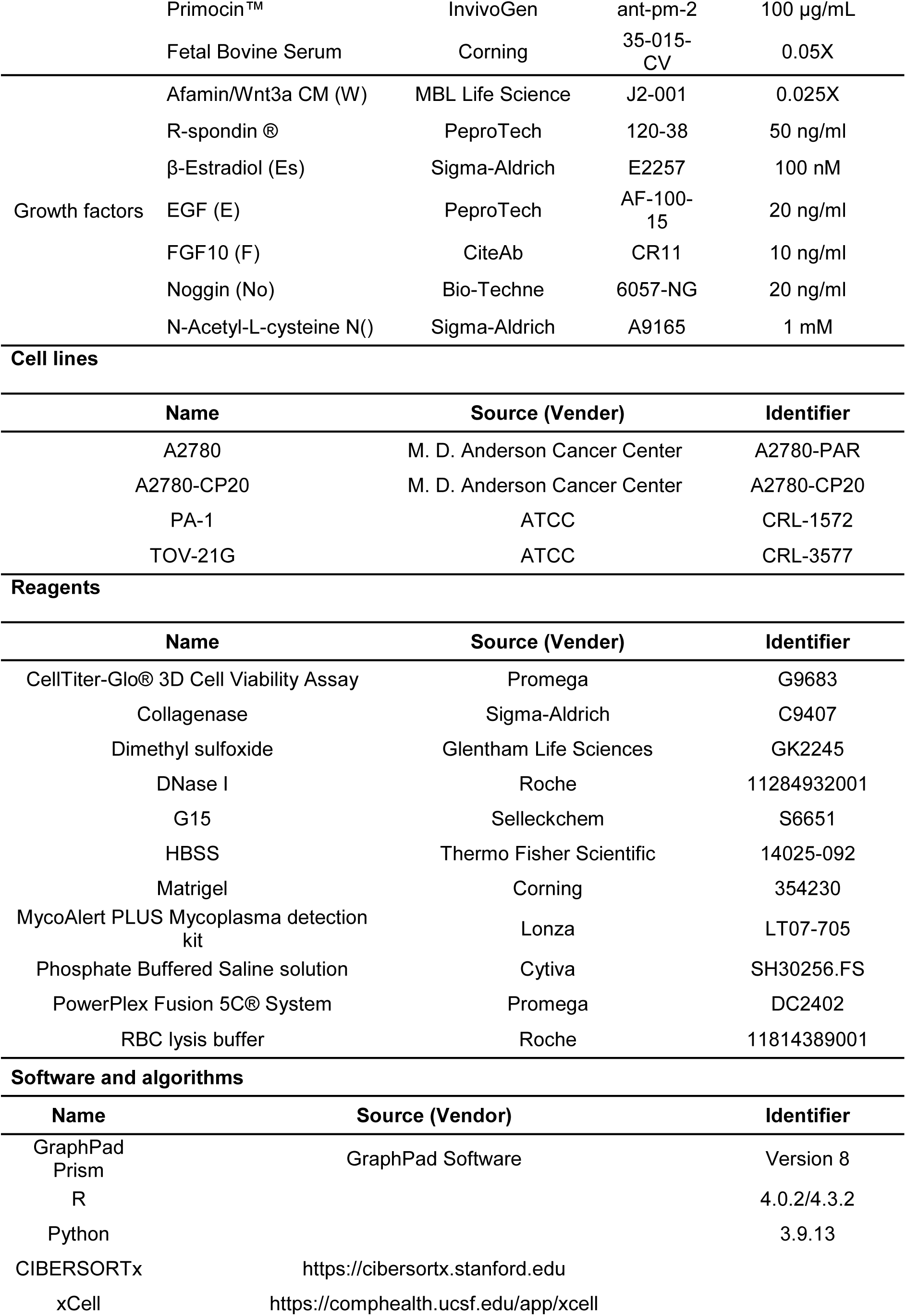
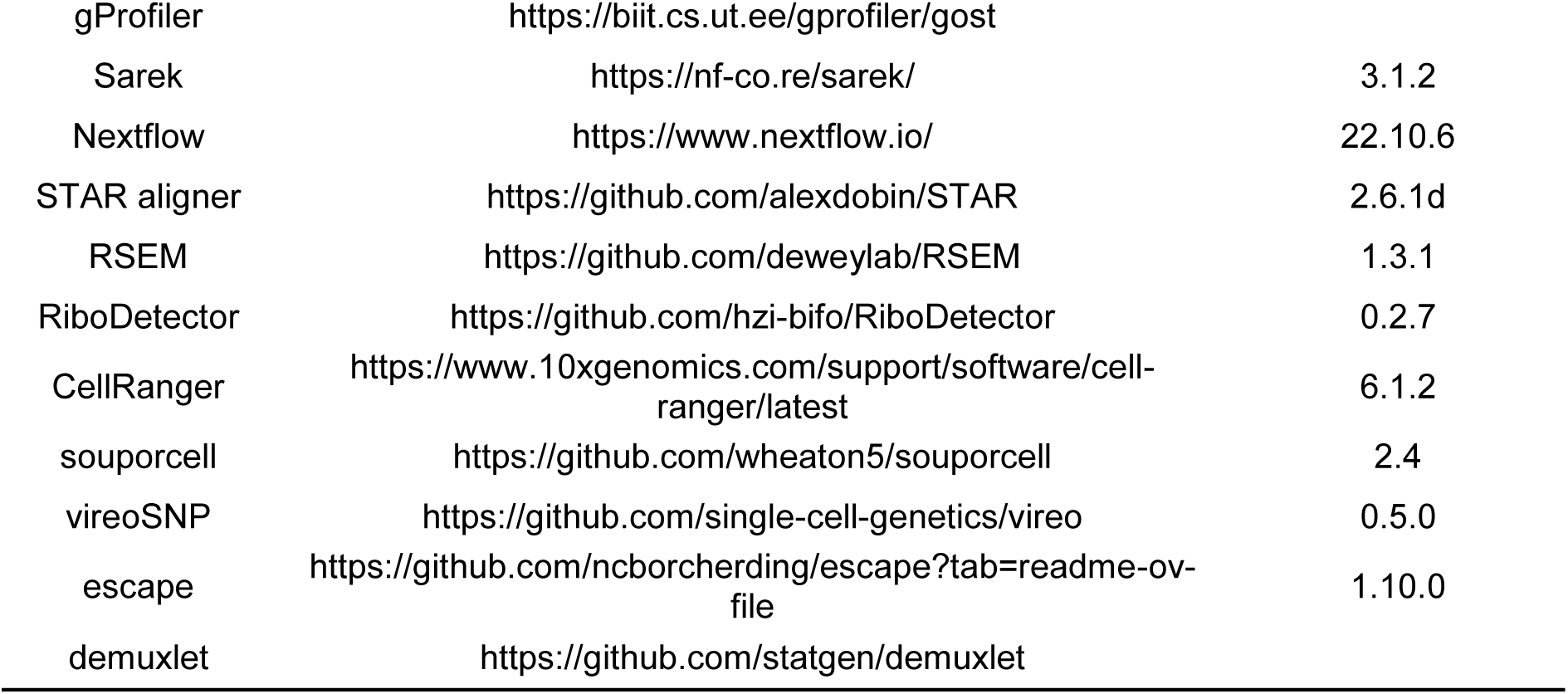

### 2.2. Biofabrication of patient-derived tumor models

Tissue Collection and Cell Culture After obtaining informed consent, tumor specimens and clinical records (Supplementary Table 1) were collected from patients undergoing surgery at Samsung Medical Center (SMC) and Seoul National University Hospital (SNUH) under approved Institutional Review Board protocols (IRB file numbers SMC 2019-07-163-019 and H-2011-127-1173). Tissue samples were snap-frozen in liquid nitrogen for genomic analysis or processed for organoid generation following established protocols [18]. Fresh tissue samples were mechanically dissociated and enzymatically digested with collagenase (Sigma-Aldrich) at 37°C under agitation for up to 30 min. When necessary, red blood cells (RBCs) were eliminated using RBC lysis buffer (Roche). Patient-derived CAFs [19] were established from the tissue dissociates of samples O010, O019, O032, and O044 by seeding 5 × 10^5^ cells per well in the assay media (Supplementary Table 2).

### 2.3. Systemic growth factor (GF) profiling

GFs (Supplementary Table 3) were dispensed into 384-well plates (761001; Nest) using an I.DOT Liquid Handler (Dispendix) with 40 µL of basal medium (M0, Supplementary Table 2) per well. Cells suspended in 50% Matrigel (Corning) were printed onto 384-pillar plates (Cellvitro, MBD) in a temperature-controlled chamber (4-8°C) to prevent premature Matrigel solidification. After 30 min of gelation at 37°C, the pillar plate was combined with the GF assay plate and cultured at 37°C with 5% CO_2_ for 7 days. Cell viability was assessed using CellTiter-Glo 3D reagent (Promega) 36, and growth rates were calculated as the ratio of day 7 to day 0 luminescence signals.

### 2.4. Data analysis

The cell proliferation measurements for the 128 GF combinations were performed in quadruplicate. Quality control was implemented using two criteria: (i) outlier removal using the 1-standard deviation method, excluding values outside (mean ± 1 standard deviation) of four-replicate data, and (ii) filtering of combinations with a coefficient of variation of ≥30% after outlier removal. Growth rates were calculated relative to day 0 measurements and converted to z-scores for normalization. Hierarchical clustering of samples based on growth scores was performed using the ComplexHeatMap R package. Detailed reagent information is provided in the Supplementary Table 11.

### 2.5. DNA/RNA isolation and next-generation sequencing analysis

DNA and RNA were extracted from 17 tissue samples using an AllPrep DNA/RNA Mini Kit (Qiagen) or a DNeasy Blood & Tissue Kit and RNeasy Mini Kit (Qiagen), according to the manufacturer’s instructions. Whole-exome sequencing libraries were prepared using an Enzymatic Fragmentation Kit v1.0, Mechanical Fragmentation Kit (Twist Bioscience), or SureSelectXT Low Input Target Enrichment Kit (Agilent). RNA-seq libraries were prepared using the Illumina RNA Prep with Enrichment (Illumina) or TruSeq RNA Access (Illumina) library preparation kits. For RNA-seq library preparation for cell line experiments, a TruSeq Stranded mRNA Sample Prep Kit (Illumina) was used.

### 2.6. Variant calling from whole-exome sequencing

Germline and somatic variant calling were conducted using the nf-core/Sarek 3.1.2 pipeline [20]. Briefly, raw sequencing data in FASTQ files were aligned to the human GRCh38 reference genome using BWA-MEM [21]. Following alignment, duplicate reads were marked using GATK Mark Duplicate Spark. The base quality scores were recalibrated using GATK BaseRecalibrator and GATK ApplyBQSR. Somatic variants, including single-nucleotide variants and small insertions/deletions, were identified using Mutect2 [22]. Germline variants were identified using Strelka2 [23]. Single-nucleotide variants and insertions/deletions were annotated using the Ensembl Variant Effect Predictor [24]. Finally, MAF files were generated using VCF2MAF and oncoplots were created using the R package maftools.

### 2.7. Bulk RNA-seq

RiboDetector was used to remove rRNA sequences from the FASTQ files [25]. Subsequently, the STAR aligner [26] was used to map the RNA-seq reads to the human reference genome (GRCh38). Read counts were processed to obtain the expected counts using the RSEM package [27]. Batch effects arising from the two RNA-seq platforms were removed using ComBat-Seq in the R package sva [28]. Finally, the normalized expression values (TPM) were calculated from the expected batch effect removed counts.

### 2.8. GF responsiveness genomic interaction analysis

The GF responsiveness for each GF–sample pair was assessed as the difference between the mean growth scores in the presence and absence of GF: responsiveness (ΔG) = mean growth score of the sample (with GF) − mean growth score of the sample (without GF). The significance of the interaction between GF responsiveness and mutation signatures was assessed using the Mann–Whitney U test, and that of the interaction between GF responsiveness and gene expression was measured using the Spearman correlation coefficient.

### 2.9. TCGA cohort analysis

We used the TCGA-OV (Ovarian Serous Cystadenocarcinoma; n = 421) dataset for analysis. Gene expression data (STAR-Counts) were downloaded using the R package TCGAbiolinks and clinical data were obtained from cBioPortal. To predict the C1/C2 subtypes of TCGA patients with ovarian cancer, we performed k-means clustering (k = 2) using the Python package scikit-learn with 150 DEGs from our 18 samples as input. The gene expression values of the TCGA samples were normalized using variance-stabilizing transformation in the R package DESeq2. For the two clusters resulting from k-means clustering, we mapped the cluster enriched in ECM and developmental functions to C1, and the cluster enriched in immune functions to C2. Functional enrichment of DEGs in TCGA patients was performed using the gProfiler [29]. The epigenetically regulated RNA expression-based (EREG.EXPss) score was developed by Malta *et al.* [30]. was used as the stemness score for the TCGA samples. The C1 and C2 scores were calculated using GSVA for the C1 and C2 upregulated DEGs.

### 2.10. Analysis of GPER signaling in human ovarian cancer cell lines

Four human ovarian cancer cell lines were used: A2780, A2780/CP20, PA-1, and TOG-21G cells (Supplementary Table 11). The cells were cultured in a cell culture medium (Supplementary Table 2). The authenticity of each cell line was verified using the PowerPlex Fusion 5C® System (Promega). The absence of mycoplasma was confirmed using the MycoAlert PLUS Mycoplasma Detection Kit (Lonza), following the manufacturer’s instructions. For GPER signaling analysis, one day after seeding, the cells were treated with the GPER inhibitor G15 (Selleckchem) at a concentration of 50 μM for 12 h. Detailed information on the reagents used is provided in the Supplementary Table 11.

### 2.11. Cell deconvolution analysis

Three cell-type ESs (ImmunoScore, StromaScore, and MicroenvironmentScore) were calculated based on TPM values using the xCell web service (http://xCell.ucsf.edu/) [31]. CIBERSORTx was used for the cell-type deconvolution of TCGA samples [32]. First, we created a signature matrix of cell subtypes in our scRNA-seq data using the ‘Create Signature Matrix’ mode. As an input, we used the normalized gene expression values of 200 down sampled cells for each subtype. Next, we calculated the cell fractions in each sample using the ‘Impute Cell Fractions’ mode with the TPM values of the TCGA samples.

### 2.12. Subtype analysis of TCGA patients

The consensus molecular subtypes of TCGA patients were determined using the R package consensusOV, which uses a random forest classifier trained on consistently subtyped tumors from multiple datasets [33]. We used the TPM value of each gene as the input and set the other options as defaults. Three tumor immune phenotypes were obtained from Desbois *et al.* [34].

### 2.13. Gene Set Enrichment Analysis (GSEA) in TCGA

R package GSVA was used to calculate the ESs of the GSEA gene sets. The input was the raw count value of each gene, and the Poisson function was used as the kernel function (kcdf = Poisson) with the other options set as the default. The curated gene sets for the four reactome pathways were downloaded from MSigDB [35]: GPER1 signaling (R-HSA-9634597), ESR-mediated signaling (R-HSA-8939211), Wnt signaling (R-HSA-195721), and Wnt signaling in cancer (R-HSA-4791275).

### 2.14. Single-cell preparation and scRNA-seq

Cryopreserved patient-derived organoids were thawed and cultured in a 50% Matrigel dome in a medium with or without estradiol for 3 days. Sequencing data were processed using CellRanger and demultiplexed based on major voting in souporcell [36], vireoSNP [37], and demuxlet [38]. The resulting barcode matrix of 24,508 cells was processed using Seurat 5.0.0 [39], which is a toolkit for scRNA-seq data analysis. Low-quality cells were excluded based on the following criteria: (i) the number of expressed genes < 250; (ii) number of expressed genes ≥ 6000; (iii) RNA count ≤ 500; (iv) mitochondrial transcripts (indicative of apoptosis) ≥ 20%; and (v) log10 (genes per unique molecular identifier) ≤ 0.8. Unique molecular identifier count data were normalized using log transformation. In total, 20 and 309 single cells were retained, respectively (Supplementary Table 7). The top 2000 highly variable genes were selected to aggregate samples into a merged dataset. The merged cell-by-gene matrix was processed using SCTransform (v2) [40] in the Seurat package, which included functions for the normalization, regression, and identification of variable genes. Principal component analysis was applied to the processed matrix. Next, we applied harmony integration [41], which effectively reduced technical batch effects while preserving biological variation. The RunHarmony function returns a Seurat object updated with the corrected harmony coordinates. Subsequently, the main cell clusters were identified using Seurat’s FindClusters function with the following options: reduction = harmony and resolution = 0.35, and visualized using Uniform Manifold Approximation and Projection.

### 2.15. PRIMUS cell subtype analysis

We applied PRIMUS [42] to identify the phenotypic cell subtypes of malignant cells, fibroblasts, and macrophages, while accounting for patient-specific components and technical noise. To determine the optimal number of clusters, we ran PRIMUS for k values ranging from 1 to 15, each with five random initial parameter sets. Based on the Bayesian information criterion, we selected k = 3, 6, and 2 for the malignant cells, fibroblasts, and macrophages, respectively.

Fibroblasts expressing both epithelial and fibroblast markers were excluded from this study. Representative gene sets were identified using the methods reported by Zhang *et al.* [43]. through the following steps. (i) Marker genes for each cluster were identified using differential expression analysis by comparing query clusters with other clusters. Genes with a log2(fold change) ≥ 0.05 and FDR < 0.05 were selected from each comparison. (ii) Using CS-core [44], a co-expression matrix was calculated to identify significantly co-expressed genes (FDR < 0.05). (iii) From the co-expression matrix, co-expression networks were extracted using the WGCNA algorithm [45], and the component genes of each co-expression network were used as representative gene sets. (iv) Finally, a gene set overrepresentation analysis was conducted for the remaining genes in each representative gene set.

### 2.16. Single-cell GSEA

The ES gene set for each cell was calculated using the R package escape. C1 and C2 scores were determined as the percentile rank difference between the ESs of the C1- and C2-upregulated DEGs. The gene set ‘Oxidative Phosphorylation’ was obtained from the cancer hallmark gene sets. The gene set ‘Antigen Presentation’ and TGF-β CAF markers were sourced from Hornburg *et al.* [46]. The CAF.S1 markers were downloaded from Kieffer et al. [47].

### 2.17. Cell-cell interaction analysis

To identify and visualize cell-cell interactions among cell subtypes and identify major signaling differences between estradiol-treated and untreated cells, the R package CellChat was used according to the developer’s vignette (https://github.com/sqjin/CellChat) [48]. We split the integrated scRNA-seq data into estradiol-treated and untreated groups and ran CellChat independently. Next, we conducted a comparative analysis of the two CellChat datasets shown in the vignette ‘Comparison analysis of multiple datasets using CellChat.’

### 2.18. Survival analysis of TCGA-OV patients

Survival information for TCGA patients was obtained from the ‘Curated survival data’ file in the TCGA Ovarian Cancer dataset of UCSC Xena (https://xenabrowser.net/datapages/), as reported by Liu *et al.* [49]. Survival and Survminer R packages were used to create Kaplan– Meier curves, and the significance of survival differences was determined using the log-rank test.

### 2.19. Quantitative analysis of FB.TNFSF10 fractions and pathway enrichment

To quantify the FB.TNFSF10 cell fractions, bulk RNA-seq data were deconvoluted using CIBERSORTx, with a cell signature matrix derived from our scRNA-seq data. The enrichment of FB.TNFSF10 cell markers was assessed using ssGSEA with the GSVA package. Three marker gene sets were used to ensure robust ssGSEA results. The first set comprised markers identified by differential expression analysis, using the Wilcoxon rank-sum test in the Seurat package. The second set included markers derived from co-expression network analysis, as shown in Supplementary Figure 13B. Third set of integrated markers from both approaches. included markers derived from the co-expression network analysis. Third set of integrated markers from both approaches.

### 2.20. Pancancer analysis of TCGA patients

Transcriptomic data from 11,096 samples representing 33 cancer types were obtained from the TCGA PancanAtlas database (https://gdc.cancer.gov/about-data/publications/pancanatlas). These batch-corrected gene expression data were used to perform GSEA of TGFβ-associated Cancer-Associated Fibroblast (TGFβ.CAF) and CAF.S1 marker signatures. In addition, the pre-computed xCell scores for the TCGA pancancer samples were retrieved from the xCell web portal (http://xCell.ucsf.edu/).

### 2.21. Xenium data analysis

We obtained preprocessed 5k Xenium data of an ovarian cancer sample from the 10x Genomics database (https://www.10xgenomics.com/datasets/xenium-prime-ffpe-human-ovarian-cancer). For the fibroblast subtype prediction, cells categorized as “tumor-associated” and “stromal-associated fibroblasts” were selected. We used the DeepCC tool [50] and trained a deep-learning model using the fibroblast transcriptome from the scRNA-seq dataset.

To analyze the correlation between fibroblast subtypes and T/NK cells, we divided the Xenium image into a 7×8 grid of tiles. Fibroblast fractions were calculated from the total fibroblast count within each tile, whereas T/NK cell fractions were calculated from the total cell count per tile to prevent bias from variation in fibroblast abundance. The ssGSEA analysis for the TGFβ CAF gene set was performed using the ’escape R package’ with parameters ’method = UCell’ and ’maxRank = 5000’. For visualization of Xenium cells, we used the scanpy and squidpy Python packages.

### 2.22. ICI-treated patients’ data collection and preprocessing

We analyzed bulk RNA sequencing (RNA-seq) data from tumor samples across 16 cohorts with ICI treatment: Kim *et al.* [51]; Huang *et al.* [52]; VanAllen *et al.* [53]; Riaz *et al.*[54]; Chen *et al.* [55]; Liu *et al.* [56]; Hugo *et al.* [57]; Gide *et al.* [58]; Prat *et al.* [59]; Lee *et al.*[60]; Cho *et al.* [61]; Jung *et al.* [62]; Hwang *et al.* [63]; Miao *et al.* [64]; Phillips *et al.* [65]; and IMvigor210 [66].

For eight datasets (Gide, Hugo, Jung, Kim, Phillips, Lee, Riaz, and Cho), raw fastq files were obtained from patient samples and aligned to the GRCh38 human reference genome using STAR (version 2.7.9a). Read quantification was performed using the RSEM pipeline (version 1.3.3). For the IMvigor210 dataset, raw read counts were directly downloaded. Gene expression levels were calculated by normalizing the counts using the trimmed means of M-value (TMM) normalization implemented in the edgeR R package (version 3.32.1) [67, 68]. For the remaining datasets (Huang, VanAllen, Liu, Chen, Hwang, and Miao), we used processed gene expression levels provided by Lee *et al.* Processed expression data from the Prat dataset were retrieved from the Gene Expression Omnibus (GEO) using the GEOquery R package (version 2.58.0).

For drug response classification, patients were categorized based on the RECIST criteria: those with a partial or complete response were classified as responders (R), whereas those with stable or progressive disease were classified as non-responders (NR). The Van Allen cohort was an exception, in which the response classification followed the criteria established by Lee *et al*.

## 3. Results

### 3.1. Integrative genomics and growth factor-omics analysis

Cancer exhibits significant molecular heterogeneity among patients, resulting in diverse GF responses that can affect treatment outcomes. Understanding these different responses is crucial for developing personalized therapeutic strategies. However, systematic approaches to characterize patient-specific GF responsiveness are limited.

To address this gap, we designed a comprehensive study to identify the GFs that influence cancer growth and their associated genomic patterns. Our approach uniquely combines growth factor screening with genomic profiling to establish correlations between cancer cell responsiveness to GF combinations and underlying molecular features. This integrative strategy aimed to uncover previously unrecognized patterns of GF responsiveness and their molecular determinants (Figure 1A).

**Figure 1.**
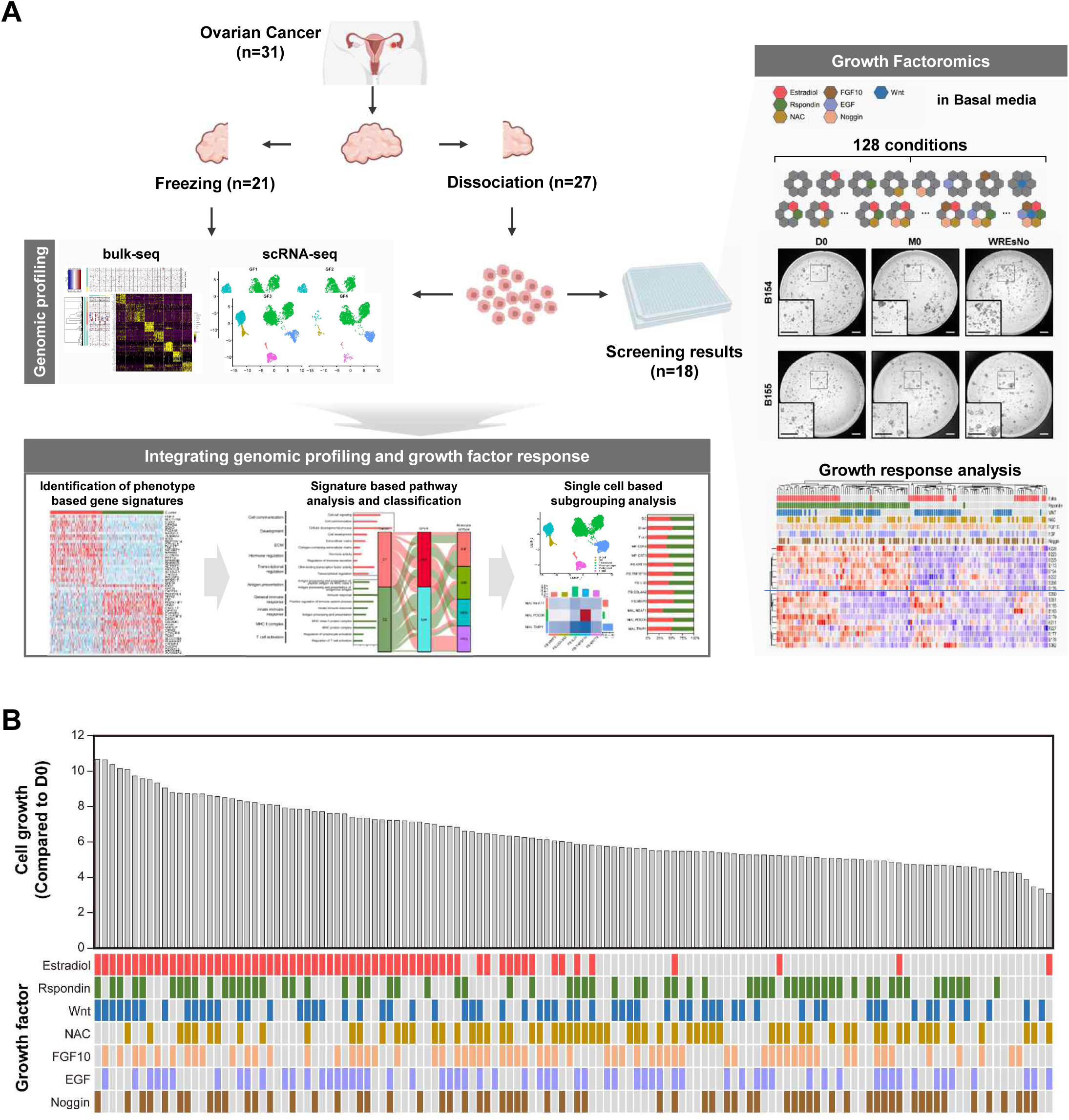
Schematic overview and growth factor-omics analysis. A. The study aims, samples, experimental scheme for growth factor-omic data acquisition, and genomics analysis are described. B. Ranking of cell growth results for 128 combinations of seven GFs. The bar plot shows the mean cell growth compared that with on day 0. The heatmap shows the combinations of the seven GFs, with gray indicating the absence of the corresponding GF.

We collected tissue samples from 31 patients (EOC) who underwent comprehensive growth factor-associated molecular profiling using multiple analytical approaches. The experimental design included parallel analyses of bulk RNA sequencing of 21 frozen specimens, and tissue dissociation of 27 samples. From the dissociated tissues, we successfully established GF profiles for 18 specimens and isolated CAFs from four samples (O010, O019, O032, and O044; Supplementary Table 1).

To investigate the effect of hormonal signaling, which is increasingly recognized as a critical modulator of tumor behavior, we performed single-cell RNA sequencing (scRNA-seq) on four dissociated tissue samples under estradiol-positive and estradiol-negative conditions. This approach allowed us to examine the cellular heterogeneity and hormone-responsive populations at an unprecedented resolution. Analysis of the successfully processed samples (n = 23) revealed a predominance of high-grade serous carcinomas (n = 19), whereas the remaining samples comprised single cases of clear cell, mucinous, endometrioid, and low-grade serous subtypes (Supplementary Table 1).

For GF screening, we developed a systematic approach using a supplemented basal medium (Supplementary Table 2) containing seven key GFs in 128 combinations (Supplementary Table 3). This comprehensive screening strategy enabled the identification of complex GF interactions that may have been missed in simpler experimental designs. Analysis of EOC-derived cell growth responses revealed a median growth rate of 5.93, with a 3.45-fold difference between the highest and lowest rates, highlighting the substantial variability in GF responsiveness among patients (Figures 1B and 2A, Supplementary Table 4).

**Figure 2.**
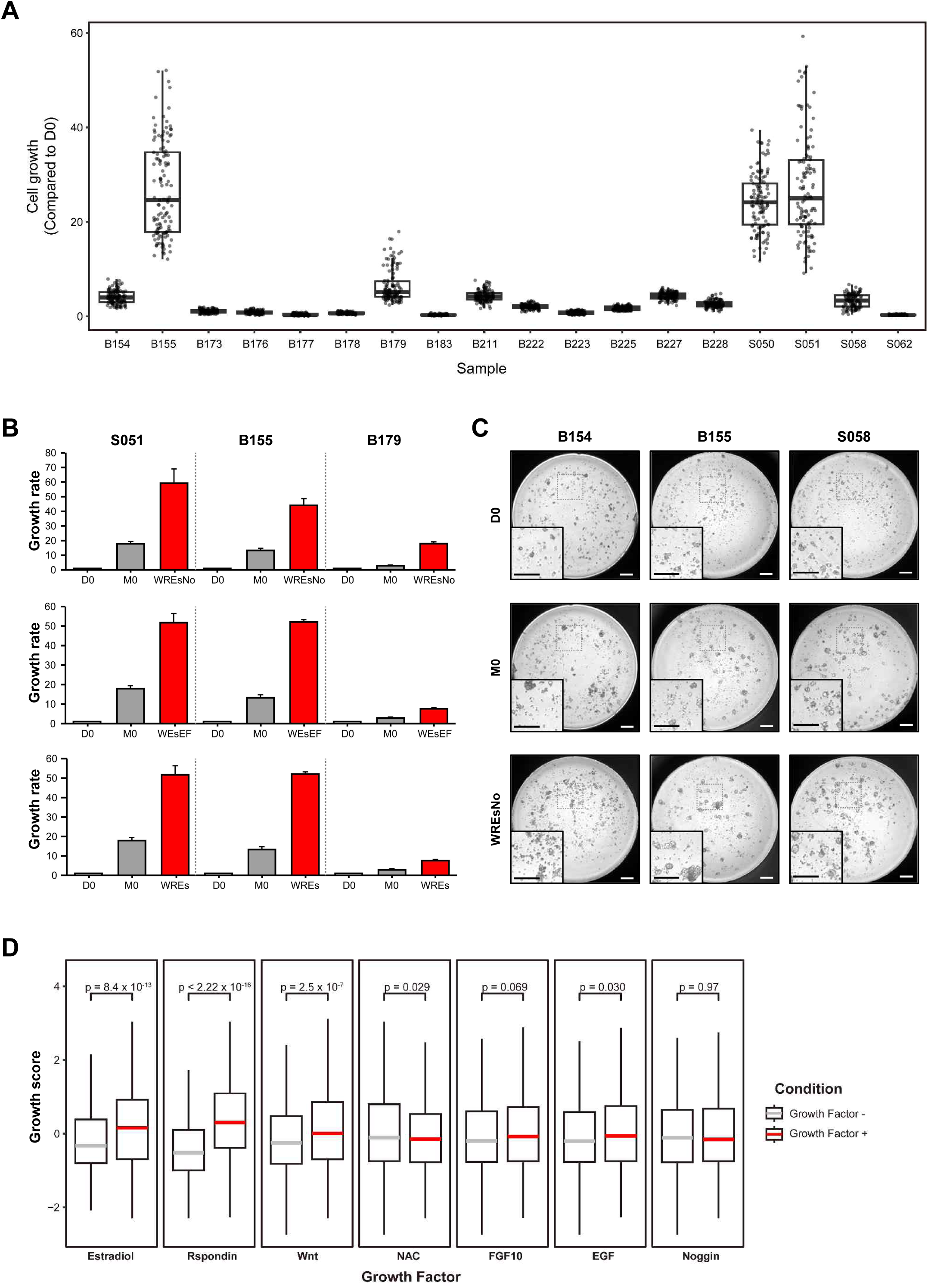
Differential cell growth induced by growth factors (GFs). A. Growth of epithelial ovarian cancer (EOC)-derived cells obtained from 18 patients after 7 days of incubation in medium containing growth factors (GFs) in 128 combinations. Box plots span the first to third quartiles and whiskers represent the 1.5 interquartile range. The central line indicates the median value. B. Cell growth efficiency of the top three growth factor-omics conditions (WREsNo (top), WEsEF (middle), and WREs (bottom)) compared to day 0 and basal medium (gray, M0). W, Wnt; R, R-spondin; Es, oestradiol; No, noggin; E, EGF; F, FGF10. C. Representative images of tumor growth under WRE sNo conditions from three independent experiments. Top: Day 0 (D0). Middle: basal medium (M0) on day 7. Bottom: WREsNo on Day 7. Scale bars, 200 μm. D. Effects of seven GFs on EOC cell growth. Cell growth was normalized for each sample using the z-score. Box plots span the first to third quartiles and whiskers represent the 1.5 interquartile range. The central line indicates the median value. The significance of the difference between the GF-positive (red line) and GF-negative (gray line) conditions was assessed using the Mann–Whitney U test.

We identified specific GF combinations that consistently promoted robust cell growth across samples. Combinations containing Wnt + R-spondin + estradiol + Noggin (WREsNo), Wnt + estradiol + EGF + FGF10 (WEsEF), and Wnt + R-spondin + estradiol (WREs) induced the highest cell growth rates (Figures 2A-C). Statistical analysis revealed that WREs significantly increased growth rate (*P <* 0.001), whereas N-acetyl-L-cysteine (NAC) reduced growth (*P =* 0.029). FGF10, EGF, and Noggin showed no significant independent impact on growth, suggesting complex interactions between GFs rather than simple additive effects (Figure 2D). These findings provide insights into the patient-specific patterns of GF responsiveness and establish a foundation for understanding how these patterns relate to underlying genomic features and potential therapeutic responses.

### 3.2. Molecular associations with GF responsiveness

Understanding the molecular basis of differential GF responses is essential for predicting treatment outcomes and for developing personalized therapeutic strategies. Although previous studies have identified individual genetic markers associated with treatment responses, a comprehensive analysis of how molecular features are related to GF responsiveness is lacking. To address these knowledge gaps, we performed hierarchical clustering analysis of standardized growth rates across our patient cohort (Figure 3A) [69]. This unbiased approach revealed two clusters, C1 and C2, suggesting differences in GF response patterns. Detailed analysis showed that C1 samples exhibited significantly higher growth rates in response to estradiol and Wnt signaling, whereas C2 samples showed higher responsiveness to R-spondin (Figure 3B). Other GFs, including NAC, FGF10, EGF, and Noggin, exhibited no significant association with either cluster, indicating specificity in the molecular determinants of the GF response (Supplementary Figure 1A).

**Figure 3.**
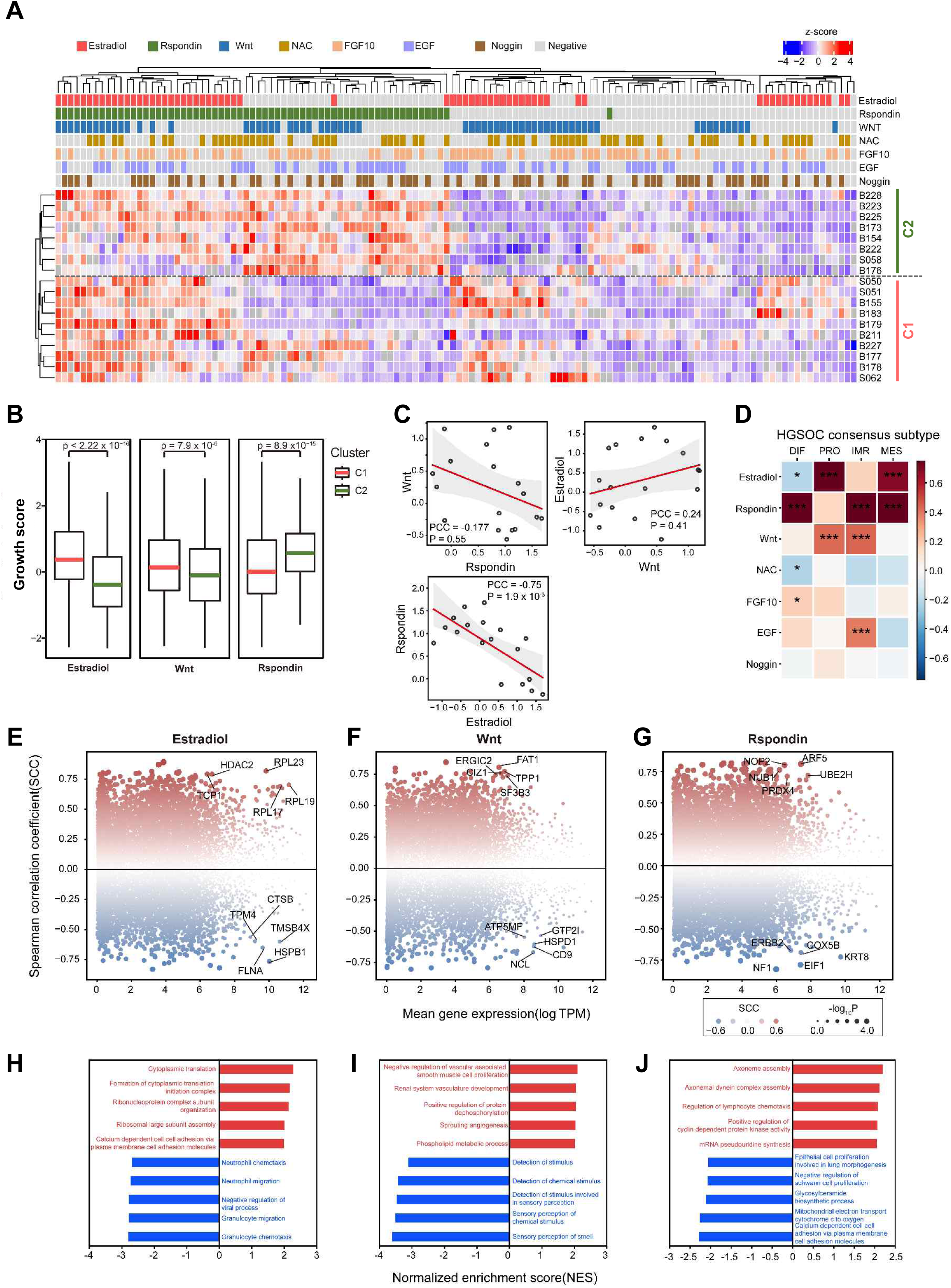
Hierarchical clustering of GF responsiveness reveals two major clusters. A. Hierarchical clustering of normalized cell growth in 18 ovarian cancer samples revealed two clusters: C1 and C2. B. Comparison of the growth scores of samples in clusters C1 (red line) and C2 (green line) cultured in the presence or absence of the indicated GFs. Box plots span from the first to third quartiles, and the whiskers represent the 1.5 interquartile range. The central line indicates the median value. *P*-values were calculated using the Mann–Whitney U test. C. Correlation of growth scores between the indicated GFs. Correlations were analyzed using PCC. D. Responsiveness of four HGSOC molecular subtypes (DIF, PRO, IMR, and MES) to the indicated GFs. The heatmap indicates the amount of change in average normalized cell growth. Significance was assessed using the Mann–Whitney U test. E, F, G. Bland–Altman (MA) plot of the correlation between gene expression and responsiveness to estradiol (E), Wnt (F), and R-spondin (G). The x-axis represents the mean gene expression across the samples and the y-axis shows the Spearman correlation coefficient (SCC). The dot size indicates the significance of the SCC. H, I, J. GSEA results of positive (red) and negative (blue) SCC between gene expression and responsiveness to estradiol (H), Wnt (I), and R-spondin (J). Gene Ontology: Biological Process gene sets were used in GSEA.

Further investigation revealed a significant inverse correlation between R-spondin and estradiol responsiveness (Pearson correlation coefficient = –0.75, *P =* 1.9 × 10⁻³; Figure 3C and Supplementary Figure 1B). This unexpected finding suggests that these pathways may have antagonistic effects, providing new insights into potential combination treatment strategies. The strong correlation also indicated these factors were the main determinants distinguishing C1 and C2 clusters.

To understand the molecular basis of these differences, we performed a differential gene expression (DEG) analysis, which identified 50 representative genes, with 25 genes specific to each cluster (Supplementary Figure 2 and Supplementary Table 5). This analysis revealed distinct molecular signatures associated with GF responsiveness. Estradiol responsiveness was significantly higher in the proliferative (PRO) and mesenchymal (MES) subtypes, whereas the differentiated (DIF) subtype showed lower estradiol and higher R-spondin responsiveness. Immunoreactive (IMR) and MES subtypes are predominantly R-spondin responsive (Figure 3D and Supplementary Figure 3) [70, 71].

Elevated estradiol responsiveness was associated with upregulation of *HDCA2* and *RPL23*, downregulation of *TMSB4X* and *HSPB1*, and enrichment of cytoplasmic translation and cell adhesion gene sets, whereas immune-related gene sets were depleted (Figures 3E and 3H). Enhanced Wnt responsiveness correlated with increased *FAT1* and *TPP1* expression, decreased *HSPD1* and *CD9* expression, and the enrichment of angiogenesis-related gene sets (Figures 3F and 3I). The heightened R-spondin response resulted in increased *ARF5* and *UBE2H* expression, reduced *KRT8* and *COX5B* expression, and enrichment of axoneme assembly and lymphocyte chemotaxis genes (Figures 3G and 3J). Integrated analysis of the mutation profiles and GF responsiveness revealed significant correlations between specific genetic alterations and GF sensitivity (Supplementary Figures 4 and 5).

### 3.3. Characteristics of C1/C2-type samples in TCGA cohort

Although our initial analysis identified distinct GF response patterns and associated molecular signatures in our patient cohort, validation using a larger independent dataset is crucial to establish the broader relevance of these findings. To validate the molecular characteristics of the C1/C2 clusters and assess their associated clinical features, we classified the samples from the TCGA-OV cohort (n = 421) into C1 or C2 using k-means clustering based on expression patterns. We identified 193 C1-type and 228 C2-type patients with no significant cluster-specific mutation patterns (Supplementary Figure 6).

Analyses of demographic and clinical characteristics revealed several significant associations. C1-type patients exhibited significantly higher age at diagnosis and aneuploidy scores than C2-type patients (*P* = 5.4 × 10⁻¹⁰ and *P* = 1.8 × 10⁻¹², respectively). However, tumor mutation burden, ESR1 expression, winter hypoxia score, MSI score, and mutation counts did not differ significantly between the groups (Supplementary Figure 7A). Overall and progression-free survival rates were similar (Supplementary Figure 7B).

DEG analysis revealed that 2,604 and 486 genes were upregulated in the C1 and C2 treatments, respectively (Figure 4A and Supplementary Table 6). C1 upregulated genes were enriched in ’collagen-containing ECM’ (FDR = 1.22 × 10⁻⁵), whereas C2 genes were enriched in ’immune response’ (FDR = 8.59 × 10⁻¹⁹, Figure 4B). The stemness score was significantly higher in C1 (*P* = 0.016; Figure 4C). C1 showed a higher StromaScore (*P* = 0.0013), whereas C2 exhibited higher ImmunoScore and MicroenvironmentScore (*P* = 7.4 × 10⁻⁹ and *P* = 4.9 × 10⁻⁷, respectively; Figure 4D).

**Figure 4.**
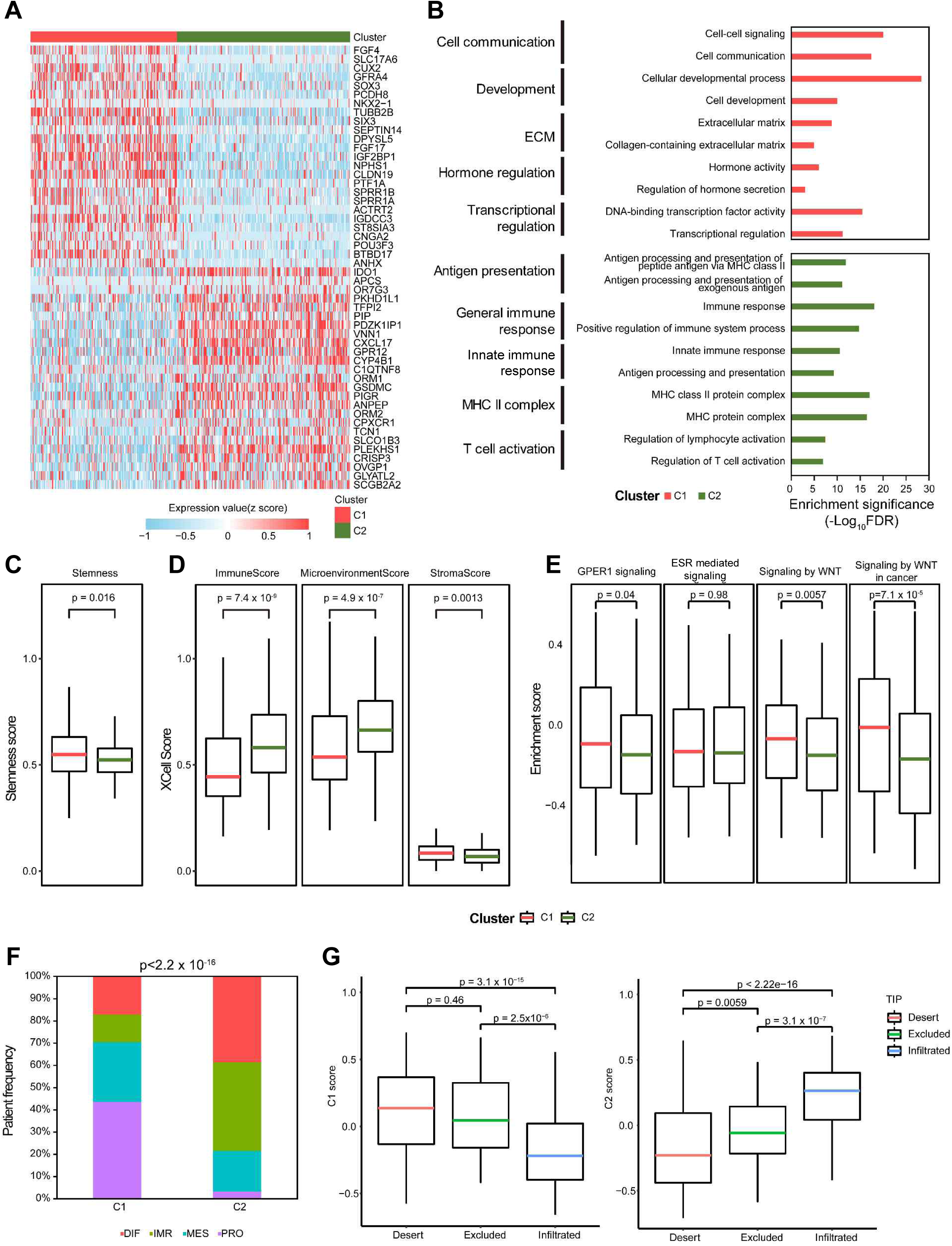
C1/C2 signature characteristics of the TCGA-OV cohort. A. Top 50 DEGs between the C1- and C2-type samples in TCGA-OV cohort. The heatmap shows the z-score-normalized gene expression of the top 25 upregulated and downregulated genes. B. GO enrichment results of DEGs between the C1-(red) and C2-type (green) samples in the TCGA-OV cohort. C. The stemness scores of the C1-type (red line) and C2-type (green line) samples were calculated using the EREG.EXPss scores. Box plots span from the first to third quartiles, and the whiskers represent the 1.5 interquartile range. The central line indicates the median value. Statistical significance was calculated using the Mann–Whitney U test. D. Cell-type ESs of C1-type (red line) and C2-type (green line) samples were calculated using xCell: ImmuneScore (composite score of immune cell types), StromaScore (composite score of stromal cell types), and MicroenvironmentScore (composite scores of immune and stromal cell types). Box plots span from the first to third quartiles, and the whiskers represent the 1.5 interquartile range. *P*-values were calculated using the Mann–Whitney U test. E. Enrichment results of C1-type (red line) and C2-type (green line) samples for four Reactome pathways: GPER1 signaling (R-HSA-9634597), ESR-mediated signaling (R-HSA-8939211), WNT signaling (R-HSA-195721), and WNT signaling in cancer (R-HSA-4791275). Box plots span from the first to third quartiles, and the whiskers represent the 1.5 interquartile range. The central line indicates the median value. *P*-values were calculated using the Mann–Whitney U test. F. Stacked bar plot displaying the distribution of HGSOC consensus subtypes in C1- and C2-type samples. Statistical significance was calculated using the chi-square test. G. ESs of immune phenotypes (tumor immune phenotype [TIP]: desert, excluded, or infiltrated) of TCGA patients in the C1-type (left) and C2-type (right) samples. Box plots span from the first to third quartiles, and the whiskers represent the 1.5 interquartile range. The central line indicates the median value. *P*-values were calculated using the Mann–Whitney U test.

Gene set variation analysis revealed significantly higher enrichment scores for G-protein-coupled estrogen receptor (GPER) signaling and Wnt-related signaling in C1 (*P* = 0.04, *P* = 0.0057, and *P* = 7.1 × 10⁻⁵, respectively; Figure 4E). ESR-mediated signaling showed no significant difference (*P* = 0.98), suggesting that GPER may be a key mediator of estradiol responsiveness [72–74].

C2 samples showed higher fractions of differentiated (DIF) and immune-reactive (IMR) subtypes, whereas C1 had higher fractions of mesenchymal (MES) and proliferative (PRO) subtypes (Figure 4F and Supplementary Figure 7C). C1 samples exhibited significantly higher deserts and excluded immune phenotypes, whereas C2 samples showed a higher infiltration phenotype (Figure 4G) [75].

These findings validate our initial observations and provide new insights into the biological and clinical significance of the C1/C2 classification. The distinct molecular and cellular characteristics associated with each type of cancer suggest potential therapeutic implications, particularly regarding immunotherapy responses and targeted intervention strategies.

### 3.4. GPER1 signaling activity correlates with MES and DIF subtypes

Understanding the molecular mechanisms that determine ovarian cancer subtypes is crucial for developing effective targeted therapies. Previous studies have demonstrated the importance of estrogen signaling in ovarian cancer progression; [76] however, the specific pathways and receptors that mediate subtype-specific effects remain poorly understood.

We investigated the factors determining the four consensus subtypes [33], focusing on estrogen responsiveness, which is crucial for the C1/C2 classification. Analysis of estrogen receptor (ESR) activity, which mediates estrogen responsiveness, did not refine subtype classification based on the C1/C2 type nor did it show differences in molecular subtypes (Supplementary Figure 8).

We then analyzed the activity of GPER, which is highly expressed in the C1 cluster. GPER-high and low subtypes were defined based on GPER1 signaling enrichment scores (ES) above or below the threshold of -0.09890369, respectively. C1-GPER-high samples were enriched in the mesenchymal (MES) subtype, C1-GPER-low samples were enriched in proliferative (PRO), C2-GPER-high samples were enriched in immunoreactive (IMR), and C2-GPER-low samples were enriched in differentiated (DIF) subtype (Supplementary Figures 9A and 4B). Similar patterns were observed in the GSE51088 and GSE26193 ovarian cancer cohorts (Supplementary Figure 10).

GPER activity significantly correlated with molecular subtypes: low GPER activity associated with higher DIF scores, whereas high GPER activity associated with higher MES scores (*P* = 1.5 × 10⁻⁹ and *P* = 4.9 × 10⁻¹⁰, respectively; Supplementary Figure 9C). Functional enrichment analysis revealed that samples with high GPER activity had higher scores for RNA processing and stem cell-related genes (e.g., FOXM1 and Oct4), whereas samples with low GPER activity had higher scores for extracellular matrix organization and immune system processes (Supplementary Figure 9D).

Experimental validation using four ovarian cancer cell lines revealed no significant difference in GPER1 expression. However, PA-1 cells exhibited markedly high GPER1 signaling activity (Supplementary Figure 9E). PA-1 cells with high GPER1 activity had a lower DIF subtype score but a significantly higher MES score than other cell lines (Supplementary Figure 9F). The inhibition of GPER1 by G15 in PA-1 cells decreased GPER1 activity, increased DIF score, and decreased MES score (Supplementary Figure 9G). These findings suggest that GPER1 activation may drive the MES subtype [74].

These results reveal a previously unrecognized role of GPER1 signaling in determining the molecular subtypes of ovarian cancer, particularly in driving the mesenchymal phenotype. The identification of GPER1 as a key regulator of subtype switching has important therapeutic implications, suggesting that targeting GPER1 signaling may provide a novel strategy for modulating the tumor phenotype.

### 3.5. ScRNA-seq defines the estradiol-responsive cell fraction

Although estrogen has been implicated in cancer immune evasion [77], the specific cellular mediators of estrogen-induced immunosuppression remain poorly defined. Previous studies have largely focused on bulk tissue responses or isolated cell populations, making it difficult to identify the key cellular fractions that directly mediate the immunosuppressive effects of estrogen.

To address these knowledge gaps and identify specific estradiol-responsive cell populations, we performed scRNA-seq on four dissociated samples with or without estradiol (Figure 5A and Supplementary Table 7). Among the 18,399 filtered cells, we identified six major cell types (Supplementary Figures 11A-B, and Supplementary Table 8). Each cluster expressed significantly higher levels of cell-type specific markers (Supplementary Figure 12).

**Figure 5.**
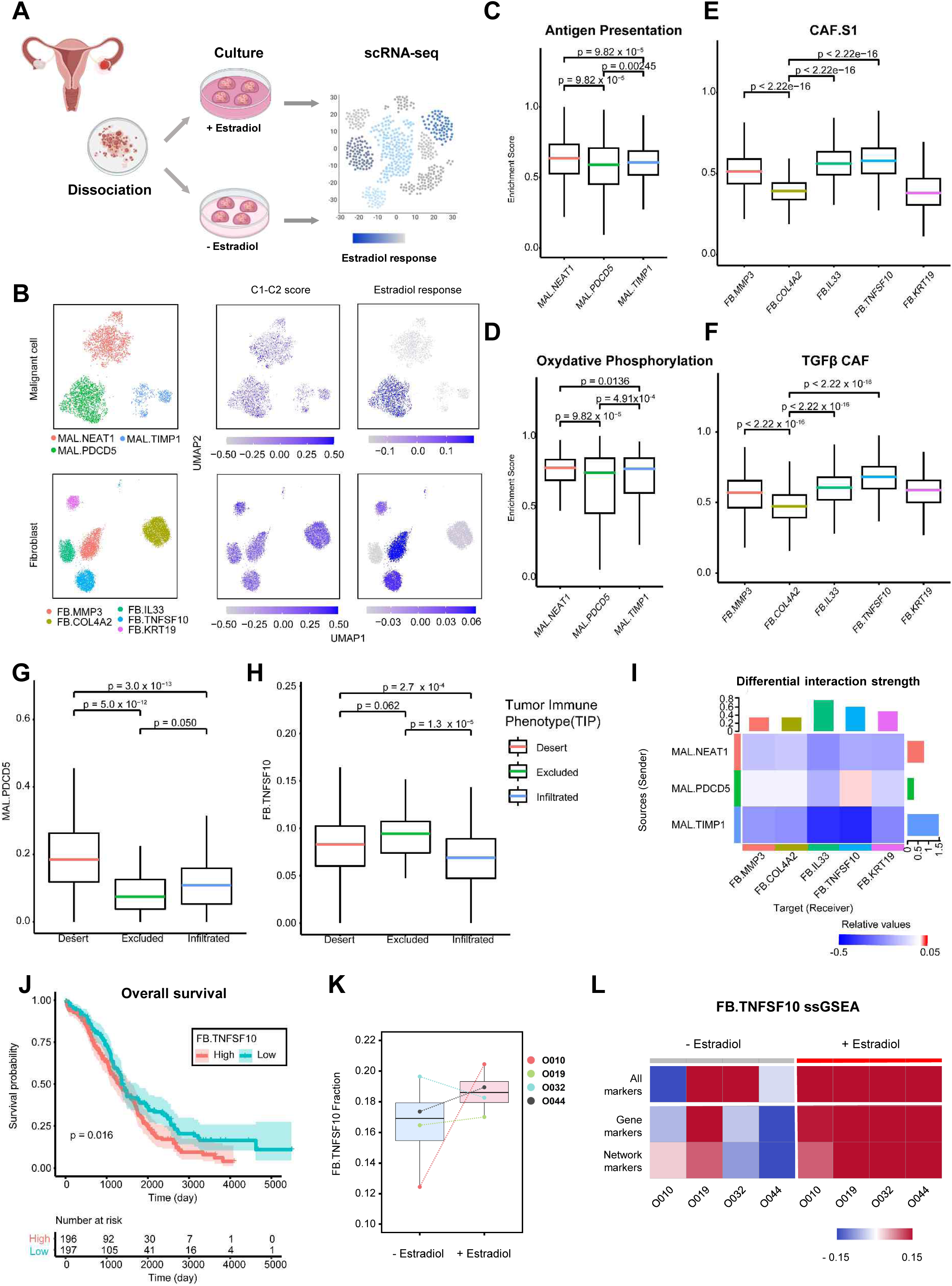
Estradiol-responsive cell fractions contribute to the immunosuppressive TME. A. A schematic representation of the scRNA-seq-based detection of estradiol-responsible cell fractions B. UMAP plot of malignant cells and fibroblast subtypes (left). C1-C2 scores based on percentile rank difference between C1 and C2 upregulated DEG ESs in subtypes (center). Estrogen response as the fold difference between estradiol-treated and untreated samples (right). C, D. Antigen presentation (C) and oxidative phosphorylation (D) ESs across malignant cell subtypes. Box plots show the quartiles and 1.5 IQR whiskers. The median is marked by a central line. Significance was determined using the Mann-Whitney U test. E, F. CAF.S1 (E) and TGF-β-CAF (F) ESs across fibroblast subtypes. Box plots show the quartiles and 1.5 IQR whiskers. The median is marked by a central line. P values from the Mann-Whitney U test. G, H. MAL.PDCD5 (G) and FB.TNFSF10 (H) cell fractions across TCGA tumor immune phenotypes were calculated using CIBERSORTx. Box plots show the quartiles and 1.5 IQR whiskers. The median is marked by a central line. Significance was assessed using the Mann-Whitney U test. I. Heatmap of the differential interaction strength between malignant cells and fibroblasts in estradiol-treated vs. untreated groups. The top and right bar plots show the total incoming and outgoing signals, respectively. Red indicates increased signaling and blue indicates decreased signaling in the treated group. J. Kaplan-Meier analysis of overall survival in FB.TNFSF10-high and -low TCGA patient groups. Subgroups were determined using CIBERSORTx, with mean FB.TNFSF10 expression as a threshold. P-values from the log-rank test. K. Box plots show FB.TNFSF10 cell fractions in the indicated patient-derived CAFs under 100 nM estradiol-treated and untreated conditions for 14 days. Box plots show the quartiles and 1.5 IQR whiskers. The median is marked by a central line. L. Heatmaps displaying ssGSEA scores for FB.TNFSF10 marker sets (All, Gene, and Network markers) in the indicated CAFs under estradiol-treated and untreated conditions.

Using PRIMUS [42], we identified three malignant (MAL), five fibroblast (FB), and two macrophage subtypes (Figure 5B, Supplementary Figures 11C-E and 13; Supplementary Tables 9 and 10) [42]. All FB populations exhibited CAF-positive patterns, with FB.KRT19 showing a relatively strong antigen-presenting CAF signature (Supplementary Figure 14) [78, 79].

The MAL.PDCD5, FB.COL4A2, FB.TNFSF10, and FB.KRT19 subtypes had higher C1 than C2 scores, and MAL.PDCD5, FB.MMP3, FB.TNFSF10, and FB.KRT19 clusters showed quantitative increases upon estradiol treatment (Figure 5B and Supplementary Figure 15). MAL.PDCD5 had a significantly lower enrichment for antigen presentation and oxidative phosphorylation functions, suggesting immunosuppressive properties (Figures 5C and 5D) [80]. Of estradiol-responsive CAFs, FB.TNFSF10 had higher scores for immunosuppressive CAF types CAF.S1 and TGF-β CAF, whereas FB.KRT19 showed lower scores (Figures 5E and 5F) [47, 81]

In TCGA samples, the desert immune phenotype [46] had a significantly larger MAL.PDCD5 fraction, whereas the excluded type had a larger FB.TNFSF10 fraction (Figures 5G and 5H). Cell-cell interaction analysis using CellChat[48] revealed that MAL.PDCD5 and FB.TNFSF10 showed the largest increase in interaction strength after estradiol treatment (Figure 5I and Supplementary Figure 16). The FB.TNFSF10-high group showed a significantly shorter overall survival (*P* = 0.016; Figure 5J).

Given the significant association between the FB.TNFSF10 fraction and patient outcomes, coupled with its distinctive tumor immune escape phenotype, we performed in-depth characterization of this cellular subset. We sought to validate whether the FB.TNFSF10 signature identified by scRNA sequencing can be translated for bulk transcriptome analysis. CIBERSORTx [32] analysis confirmed substantial CAF fractions (> 0.6) across the four established CAF cell lines, with consistent proportions maintained regardless of estradiol treatment (Supplementary Figure 17). Three out of the four CAF cells exhibited increased FB.TNFSF10 fractions following estradiol treatment, with O010 cells showing the most pronounced response (0.12 to 0.20, representing a ∼1.7-fold increase; Figure 5K). Single-sample Gene Set Enrichment Analysis (ssGSEA) of the FB.TNFSF10 gene set revealed a remarkable upregulation in response to estradiol treatment, which was consistently observed across both gene and network markers (Figure 5L).

These findings provide the first detailed cellular map of estradiol responsiveness in ovarian cancer, revealing specific cell populations that mediate hormone-dependent changes in the tumor microenvironment.

### 3.6. Estradiol-responsive cell fractions correlate with pancancer immunosuppression

Although estrogen signaling has traditionally been studied in hormone-dependent cancers, emerging evidence suggests a broader role in modulating immune responses across various cancer types [17, 77]. However, the extent to which estrogen-responsive cell populations influence the immune landscape beyond hormone-dependent cancers remains unclear.

To elucidate the relationship between estradiol and the TME, we conducted a cell deconvolution analysis of 11,069 TCGA transcriptome samples. The normalized proportions of FB.TNFSF10 and MAL.PDCD5 cells varied across the 33 TCGA cancer types (Supplementary Figure 18). We found significant negative correlations between MAL.PDCD5 and various immune cell populations, including dendritic cells, CD8+ T cells, and B cells, across the pancancer cohort (Figure 6A, left). FB.TNFSF10 exhibited positive correlations with TGFβ-CAF and CAF.S1 fractions, consistent with patterns observed in TCGA ovarian cancer cohort (Figure 6A, right) [82].

**Figure 6.**
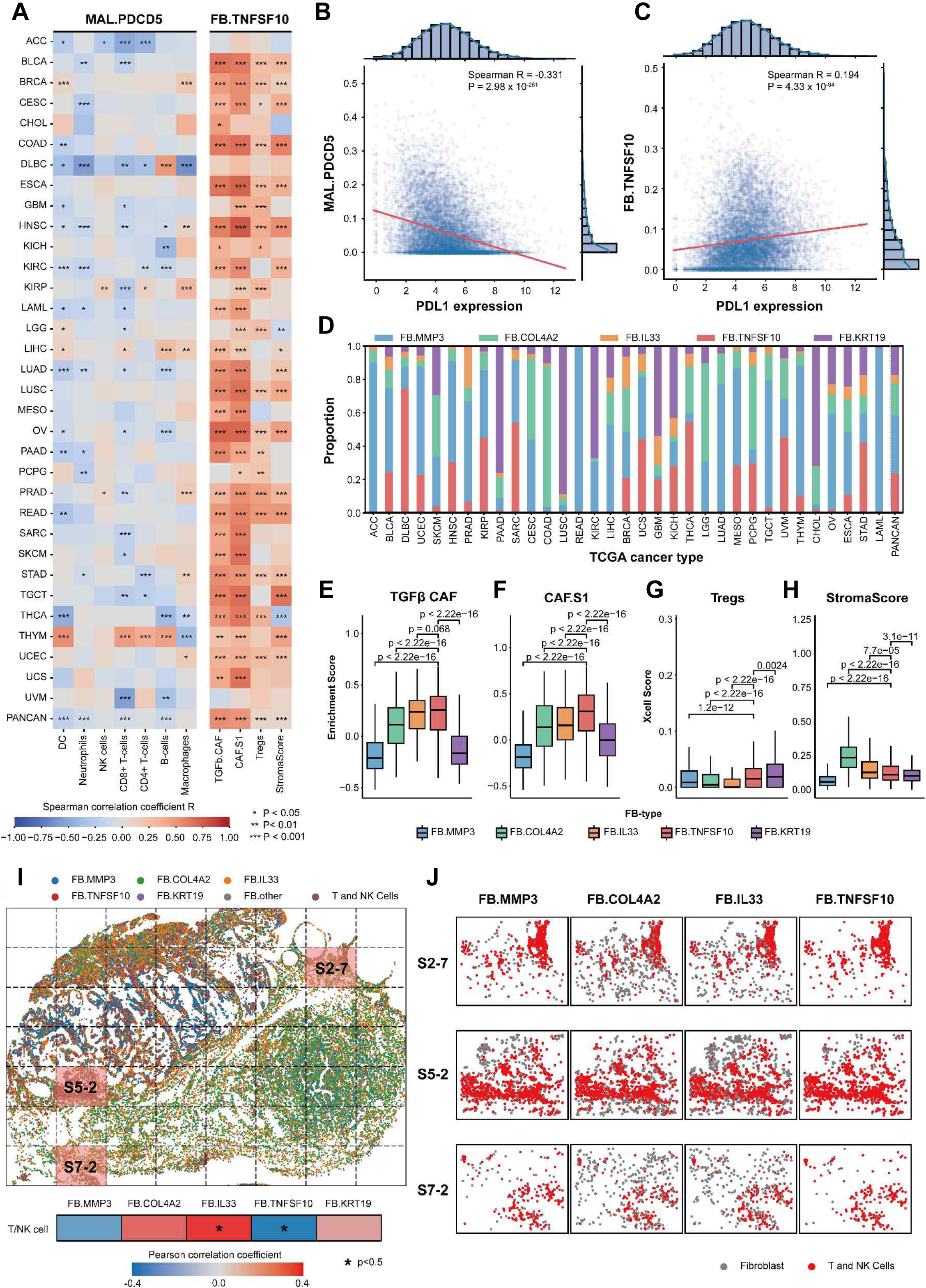
Pan-cancer immune phenotypes associated with estradiol-responsive fractions. A. A heat map of correlations between immune phenotypes and FB.TNFSF10 and MAL.PDCD5 expression across 33 TCGA cancer types. B, C. Spearman correlation analysis of PDL1 expression with FB.TNFSF10 (B) and MAL.PDCD5 (C) cell abundance in TCGA pan-cancer dataset. D. Distribution of fibroblast (FB) subtypes across the TCGA samples. FB subtype classification was based on the predominant fibroblast population identified by CIBERSORTx analysis for each patient. E, F. Enrichment scores (ESs) for TGF-β CAF (E) and CAF.S1 (F) signatures stratified by FB subtype. G, H. X-cell-derived regulatory T cell (Treg) scores (G) and stromal scores (H) across patient FB subtypes. Box plots show the median (center line), first to third quartiles (box), and 1.5× interquartile range (whiskers). Statistical significance was assessed using the Mann-Whitney U test. I. Predicted fibroblast and T/NK cell subtypes in xenium ovarian cancer samples (top). The Xenium data were divided into 56 tiles (7×8 grids, labeled S1-1 through S7-8). Pearson correlation coefficients (PCC) between normalized cell fractions of fibroblast subtypes and T/NK cells across tiles are shown (bottom). Asterisks indicate a significant correlation (*P* < 0.05). J. Distribution of four fibroblast subtypes and T/NK cells in three representative tiles (S2-7, S5-2, and S7-2), where FB.TNFSF10 cells numbered < 100. Gray dots represent fibroblast subtypes and red dots indicate T/NK cells.

Analysis of PD-L1 expression [83], revealed a significant inverse correlation with MAL.PDCD5 fraction (Spearman *R =* -0.331, *P* = 2.98 × 10⁻²⁸¹, Figure 6b). Conversely, FB.TNFSF10 demonstrated a positive correlation with PD-L1 expression (Spearman *R =* 0.194, *P* = 4.33 × 10⁻⁹⁴, Figure 6C). These findings prompted us to focus on subsequent ICI treatment analyses of FB.TNFSF10.

Next, we classified patients into five categories based on their predominant fibroblast (FB) subtype, with the distribution of these FB types varying across the 33 TCGA cancer types (Figure 6D). Consistent with our previous observations, FB.TNFSF10-type patients exhibited the highest ES for TGFβ-CAF and CAF.S1 markers among the five FB subtypes (median ES = 0.257 and 0.312, respectively; Figs. 6E and 6F) [47, 81]. Furthermore, this subtype ranked second and third in xCell scores for Tregs and stromal content, respectively (median ES = 0.0149 and 0.110, respectively; Figures 6G and 6H).

To investigate the spatial relationships between FB subtypes, we analyzed 5k Xenium data from an ovarian cancer sample (Figure 6I). We quantified the cell-type fractions across a 7×8 grid of tiles (Supplementary Figure 19A) and discovered that FB.TNFSF10 cells showed mutual exclusivity with T and NK cells (Figure 6J). Across all 56 tiles, the normalized fraction of FB.TNFSF10 cells displayed a significant negative correlation with T/NK cell fractions (PCC r= -0.317, *P* = 0.0185; Figure 6I and Supplementary Figure 19B). Consistent with previous results, FB.TNFSF10 cells exhibited elevated TGFβ CAF signature scores in the Xenium data (median ES = 0.520; Supplementary Figures 19C-D).

These findings provide compelling evidence that estrogen-mediated signaling shapes the immunosuppressive landscape across diverse cancer types through conserved cellular mechanisms. Furthermore, high-resolution spatial transcriptomic analysis revealed consistent patterns of immune cell exclusion and stromal modulation associated with FB.TNFSF10 cell population. This spatial validation provided crucial contextual evidence for the functional role of FB.TNFSF10 cells within the complex tissue architecture, highlighting their potential utility as histopathological markers.

### 3.7. Estradiol-responsive CAF fraction associates with ICI therapy resistance

Although ICIs have shown remarkable efficacy against various cancers, understanding their mechanisms of resistance remains a critical challenge. Our identification of estradiol-responsive cellular populations with immunosuppressive properties raises the possibility that these cells may influence immunotherapy outcomes. The strong association between FB.TNFSF10 and immunosuppressive features suggests its potential role in treatment resistance. In addition, as previously discussed, the MAL.PDCD5 fraction showed an inverse correlation with PD-L1 expression, which is a target of ICI therapy (Figure 6B).

We focused our subsequent analysis on the FB.TNFSF10 fraction and analyzed the distribution of FB.TNFSF10 between the responder and non-responder groups across the 18 ICI-treated cohorts. FB.TNFSF10-type patients exhibited significantly lower response rates compared to other FB-type patients (Fisher’s exact test *P =* 3.96 × 10⁻³ in the total cohort; Figure 7A). This pattern was remarkably consistent across 15 of the 18 cohorts, with FB.TNFSF10 subtypes consistently underrepresented in responders compared with non-responders (Figure 7B).

**Figure 7.**
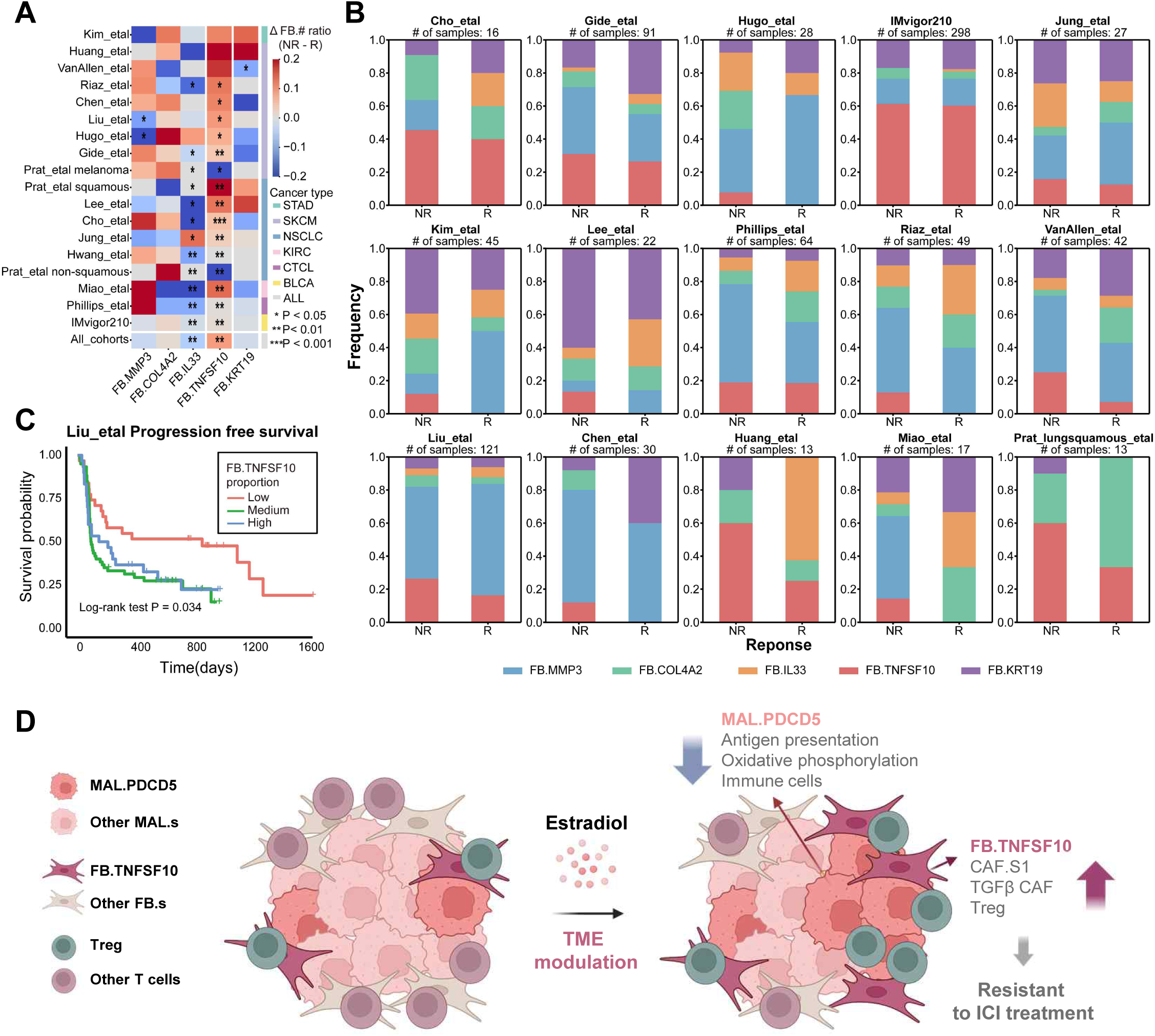
Effect of estradiol-related TME subtypes on ICI treatment response. A. Differences in FB subtype ratios between ICI responders and non-responders across the 18 cohorts. Asterisks indicate p values from Fisher’s exact test comparing the corresponding cell types between responders and non-responders. B. Comparison across 15 cohorts showing lower proportions of FB.TNFSF10-types in responders than in non-responders. C. Kaplan-Meier analysis of progression-free survival in three patient groups from the Liu et al. cohort: FB.TNFSF10-high, FB.TNFSF10-medium, and FB.TNFSF10-low. The first and third quartiles of the FB.TNFSF10 proportions were used as thresholds. The p-value was determined using the log-rank test. D. Schematic representation of estradiol-modified TME influencing ICI treatment response.

To validate these findings in terms of clinical outcomes, we analyzed progression-free survival in a cohort study by Liu *et al.* [56]. Patients with low FB.TNFSF10 expression (fourth quartile) demonstrated significantly longer progression-free survival than those with medium (second and third quartiles) or high (first quartile) expression levels (log-rank test, *p =* 0.034; Figure 7C).

These results establish that FB.TNFSF10 is a key mediator of immunotherapy resistance. In tumors exposed to estrogen, including estradiol, increased distribution of estradiol-responsive cell fractions contributes to an immunosuppressive microenvironment through two mechanisms: MAL.PDCD5-mediated inhibition of immune cell infiltration through reduced antigen presentation and oxidative phosphorylation, and FB.TNFSF10-driven immunosuppression through enhanced CAF.S1 and TGF-β CAF populations, and increased Treg infiltration (Figure 7D).

The consistency of these findings across multiple cohorts suggests a conserved mechanism of resistance, with implications for patient stratification and therapeutic intervention. These results provide a rationale for developing combination strategies that target estrogen-responsive stromal populations to enhance immunotherapy efficacy.

## 4. Discussion

Our biofabricated tumor model system addresses critical unmet needs in cancer research by revealing estradiol-responsive cellular fractions that drive immunosuppression across diverse cancer types. More importantly, this work establishes a transformative biofabrication platform that could revolutionize how we model complex tissue environments and develop therapeutic strategies.

The biofabrication approach employed in this study offers several key advantages over traditional culture systems. First, our patient-derived 3D culture platform maintains the cellular heterogeneity and architectural complexity characteristic of native tumors while enabling systematic manipulation of growth factor environments. This approach allowed us to identify specific estradiol-responsive populations that would be difficult to characterize in conventional 2D culture systems [3]. Second, our system provides a more comprehensive model of the TME, enabling investigation of tumor-stromal interactions that are critical for understanding immunotherapy resistance [16].

The discovery that FB.TNFSF10 correlates with immunotherapy resistance across multiple cancer cohorts represents a significant advancement in biomarker development. This finding is particularly noteworthy because it transcends traditional boundaries of hormone-dependent cancers, suggesting a fundamental biological principle that could revolutionize patient stratification for immunotherapy [84]. The consistency of this correlation across various malignancies indicates a conserved mechanism of estradiol-mediated immune modulation that can be effectively modeled using our biofabrication platform.

Our spatial transcriptomic analysis of engineered tissue constructs revealed compelling evidence of immune cell exclusion mediated by FB.TNFSF10 cells. This spatial validation provides crucial contextual evidence for the functional role of these cells within the complex tissue architecture, highlighting the importance of 3D tissue models in understanding cellular interactions. The mutual exclusivity between FB.TNFSF10 cells and T/NK cells observed in our biofabricated models suggests active immune exclusion mechanisms that would be difficult to identify in traditional culture systems.

Our systematic growth factor profiling approach represents a paradigm shift in biofabrication methodology that extends far beyond cancer research. The platform’s ability to systematically optimize culture conditions for patient-specific requirements makes it applicable to engineering diverse tissue constructs, from organ-on-chip models to regenerative medicine applications. The 128-combination screening approach can be adapted for any tissue type, providing a standardized framework for biofabrication across multiple research domains. This systematic approach enables the development of reproducible protocols that can be implemented across different laboratories and manufacturing facilities, addressing a critical need in the biofabrication field for consistent methods that can be translated from research to clinical applications.

The integration of 3D culture systems with comprehensive molecular profiling offers unprecedented opportunities for pharmaceutical applications and high-throughput drug screening. The principles underlying our growth factor-responsive culture systems can be adapted for engineering other complex tissue constructs that require precise control of cellular heterogeneity and signaling environments. Potential applications include modeling neurodegenerative diseases, cardiovascular disorders, and inflammatory conditions. The platform’s ability to maintain patient-specific characteristics while enabling systematic experimental manipulation provides a powerful tool for understanding tissue development and disease progression across multiple biological systems, ultimately advancing personalized medicine approaches beyond oncology.

## 5. Conclusion

The potential for translating our biofabrication platform to clinical applications is significant. The ability to rapidly generate patient-specific tumor models for drug testing could revolutionize personalized medicine approaches in oncology. Furthermore, the identification of biomarkers using our 3D culture systems could lead to more accurate predictive tests for immunotherapy responses, ultimately improving patient outcomes and reducing healthcare costs.

## Data and code availability

Raw FASTQ files of the whole-exome, bulk RNA, and scRNA sequencing data were deposited in the European Genome-Phenome Archive (EGAS50000000422) and are available from the corresponding authors upon reasonable request. TCGA gene expression data (STAR-Counts) were downloaded using the R package TCGAbiolinks, and clinical data were obtained from cBioPortal (https://www.cbioportal.org/study/summary?id=ov_tcga_pub/). The public ovarian cancer bulk RNA-seq data reanalyzed in this study are available in the Gene Expression Omnibus under accession codes GSE51088 and GSE26193. The batch-corrected transcriptomes of 11,096 samples across 33 TCGA tumor types are available in the PancanAtlas (https://gdc.cancer.gov/about-data/publications/pancanatlas).

All relevant packages and software information are provided in the Star Methods and Supplementary Table 12. No custom codes were used in this study.

## Acknowledgments

This work was supported by the Creative-Pioneering Researchers Programme in SNU; the Cooperative Research Program of Basic Medical Science and Clinical Science from SNUCM (800-20240351), the National Research Foundation (NRF) of Korea funded by the Korean government (MIST) (NRF2020M3A9D803800912, NRF2022R1A5A102641311, RS-2023-00272547, RS-2023-00222910, and RS-2023-00208519); the Ministry of Science and ICT (2024-22030007-30); the Global Physician-Scientist Training Program from the Korea Health Industry Development Institute (KHIDI, RS-2024-00440302); and the Commercialization Promotion Agency for R&D Outcomes (COMPA).

## Declaration of interests

The authors declare that they have no conflicts of interest.

## Supplementary figures and legends

**Supplementary Figure 1.**
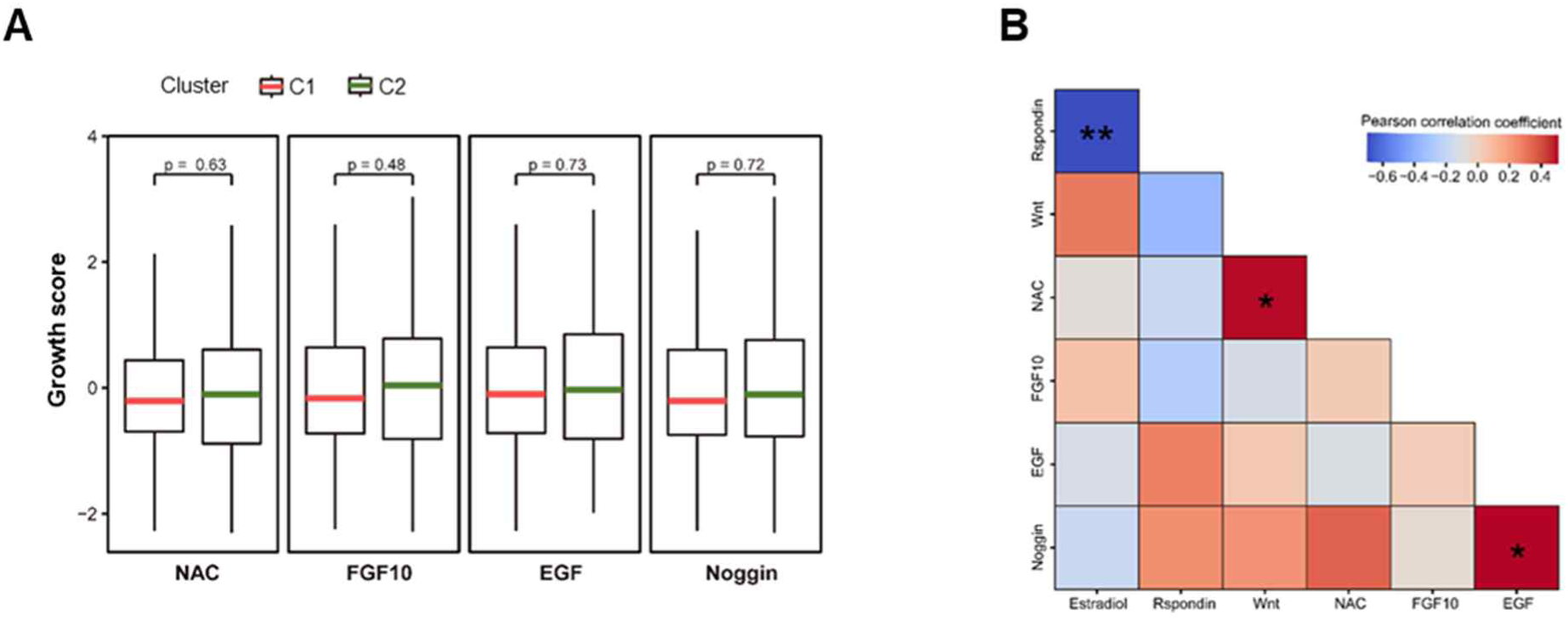
Correlations between responsiveness to the seven GFs. A. Normalized cell growth of samples from clusters C1 and C2 in the presence of NAC, FGF10, EGF, or Noggin. B. Pearson correlation coefficients between the indicated GFs.

**Supplementary Figure 2.**
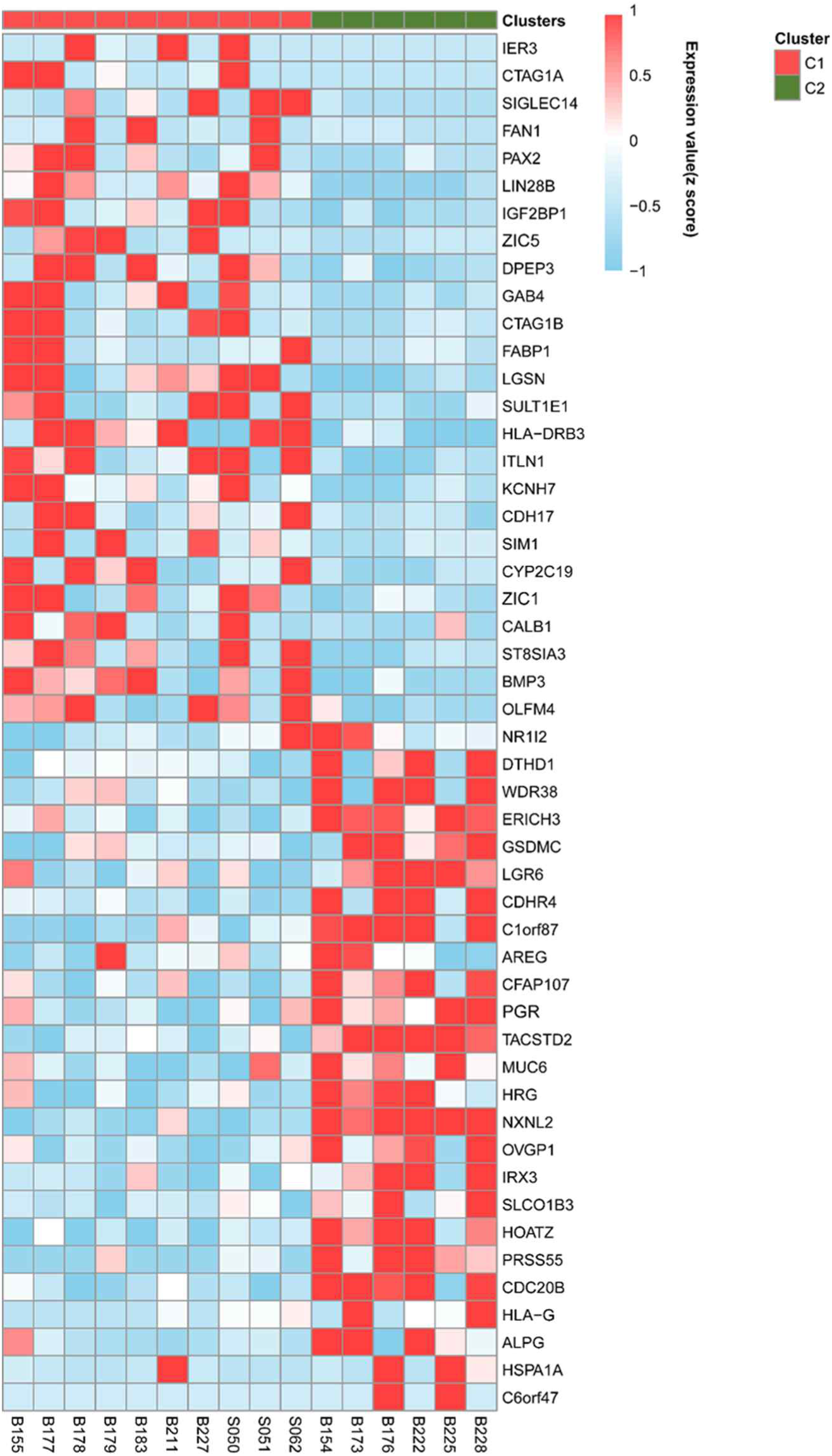
Top 50 differentially expressed genes (DEGs) used in C1/C2 clustering. The heatmap shows the z-score-normalized expression of the top 25 upregulated and downregulated DEGs.

**Supplementary Figure 3.**
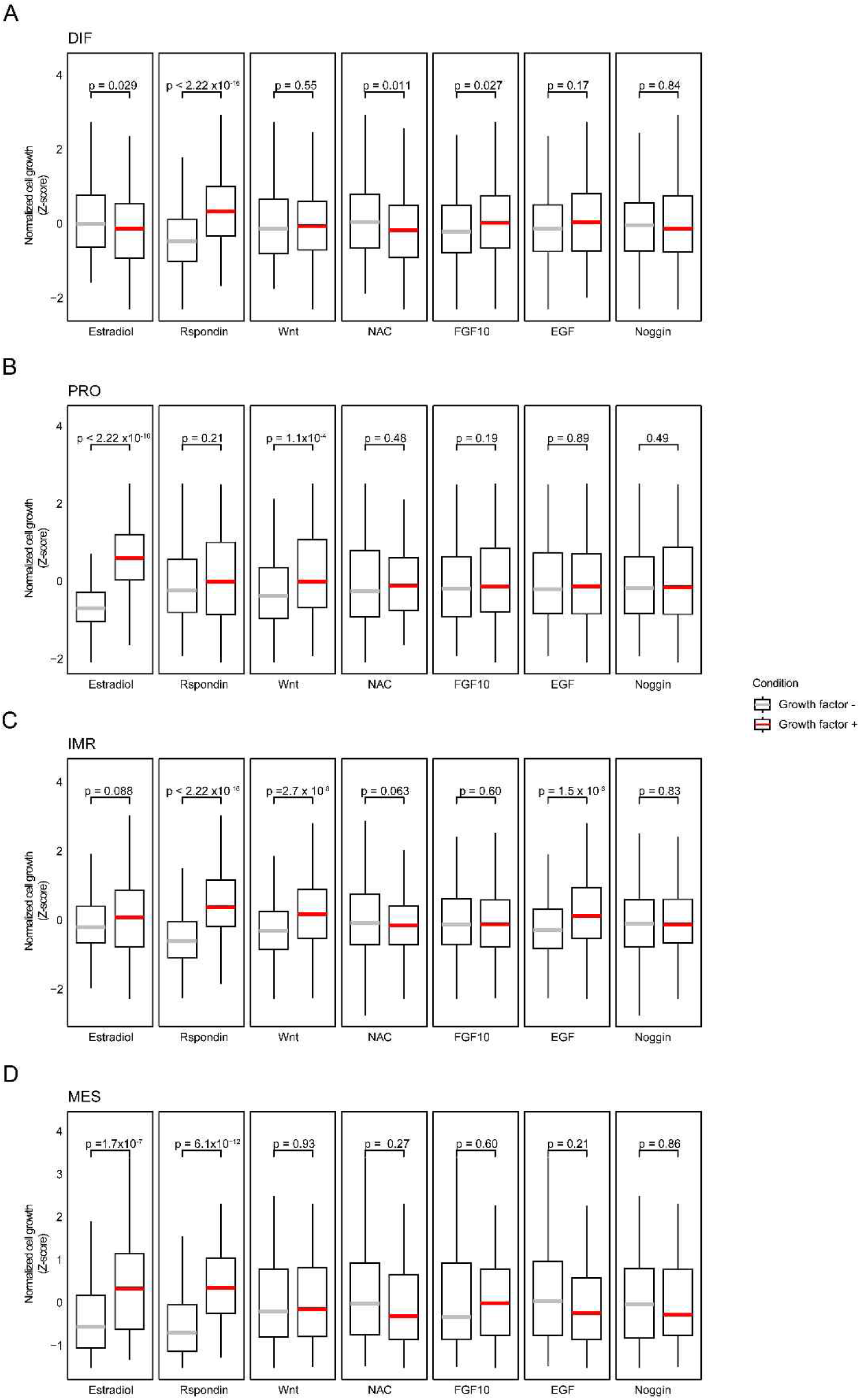
GF responsiveness of four HGSOC consensus subtypes. A–D. Growth scores of DIF (A), PRO (B), IMR (C), and MES (D) subtype samples incubated in the presence or absence of the indicated GFs. Box plots span the first to third quartiles and whiskers represent the 1.5 interquartile range. The central line indicates the median value. *P*-values were calculated using the Mann–Whitney U test.

**Supplementary Figure 4.**
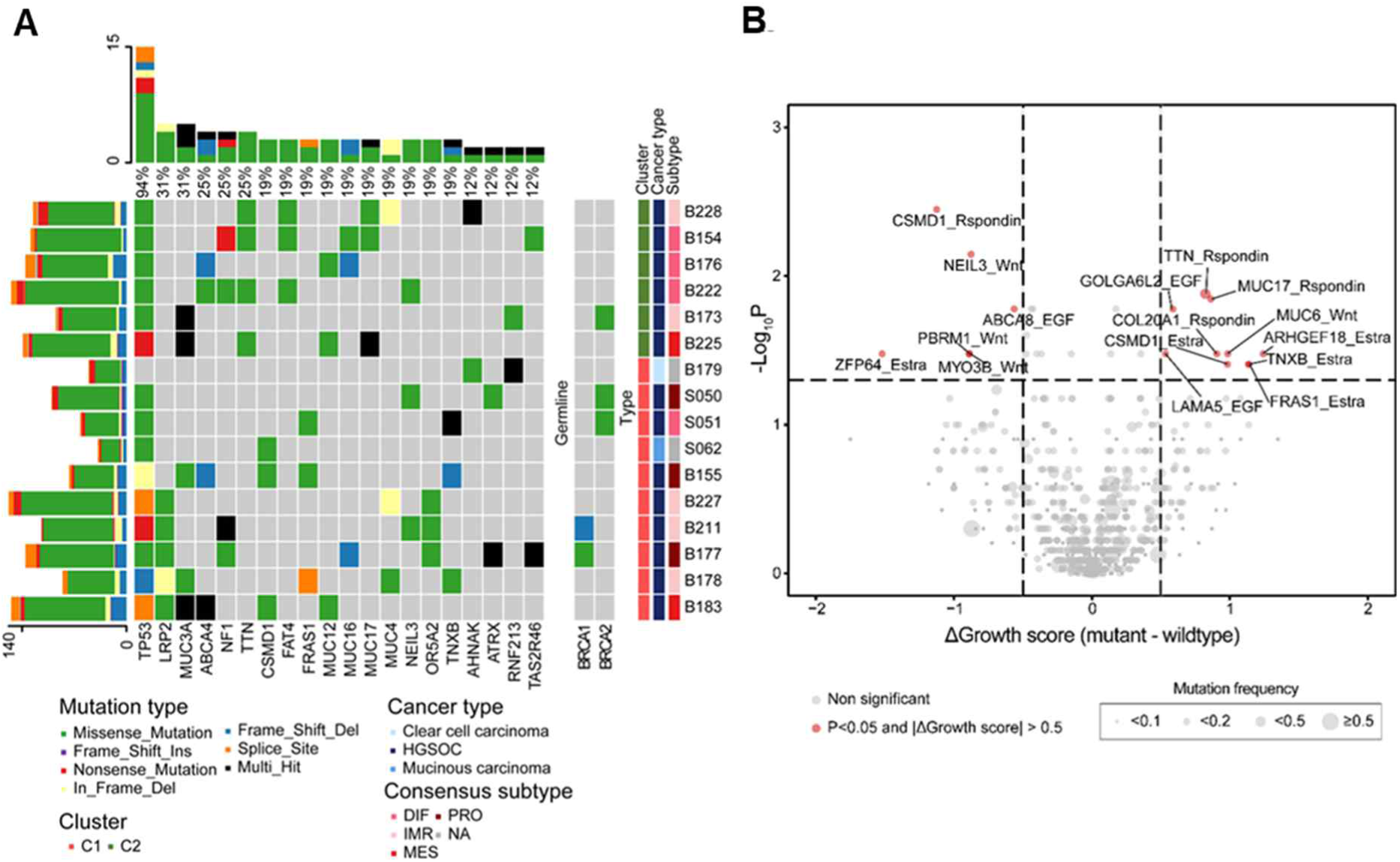
Mutation signatures associated with the GF responses. A. Oncoplot of 20 frequently mutated genes and *BRCA1/2* in the 16 EOC samples. Mutation type, C1/C2 cluster, pathological cancer type, and consensus subtype of each sample are shown. B. Volcano plot showing pairs of mutations and GFs that exhibited significant differences in growth scores between mutant and wild-type cells under the corresponding GF conditions. The x-axis represents the difference in cell growth scores, and the y-axis shows the significance of cell growth score differences, as assessed using the Mann–Whitney U test. Dot size indicates mutation frequency across samples.

**Supplementary Figure 5.**
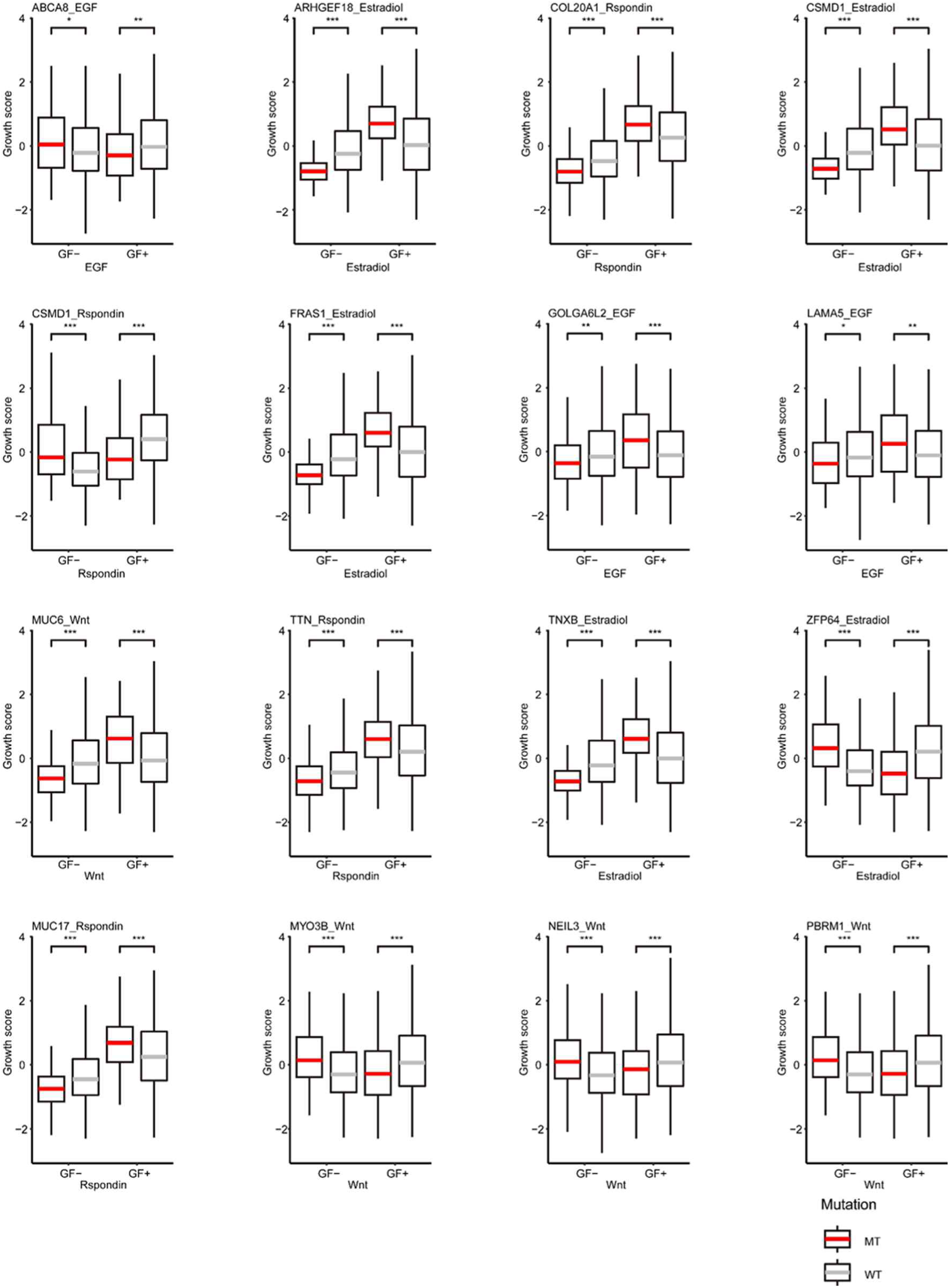
GF responsiveness according to the mutation status. Growth scores of samples harboring mutations (red line) in the indicated genes or of WT cells (gray line) cultured in the presence or absence of the indicated GFs. Box plots span the first to third quartiles and whiskers represent the 1.5 interquartile range. The central line indicates the median value. *P*-values were calculated using the Mann–Whitney U test.

**Supplementary Figure 6.**
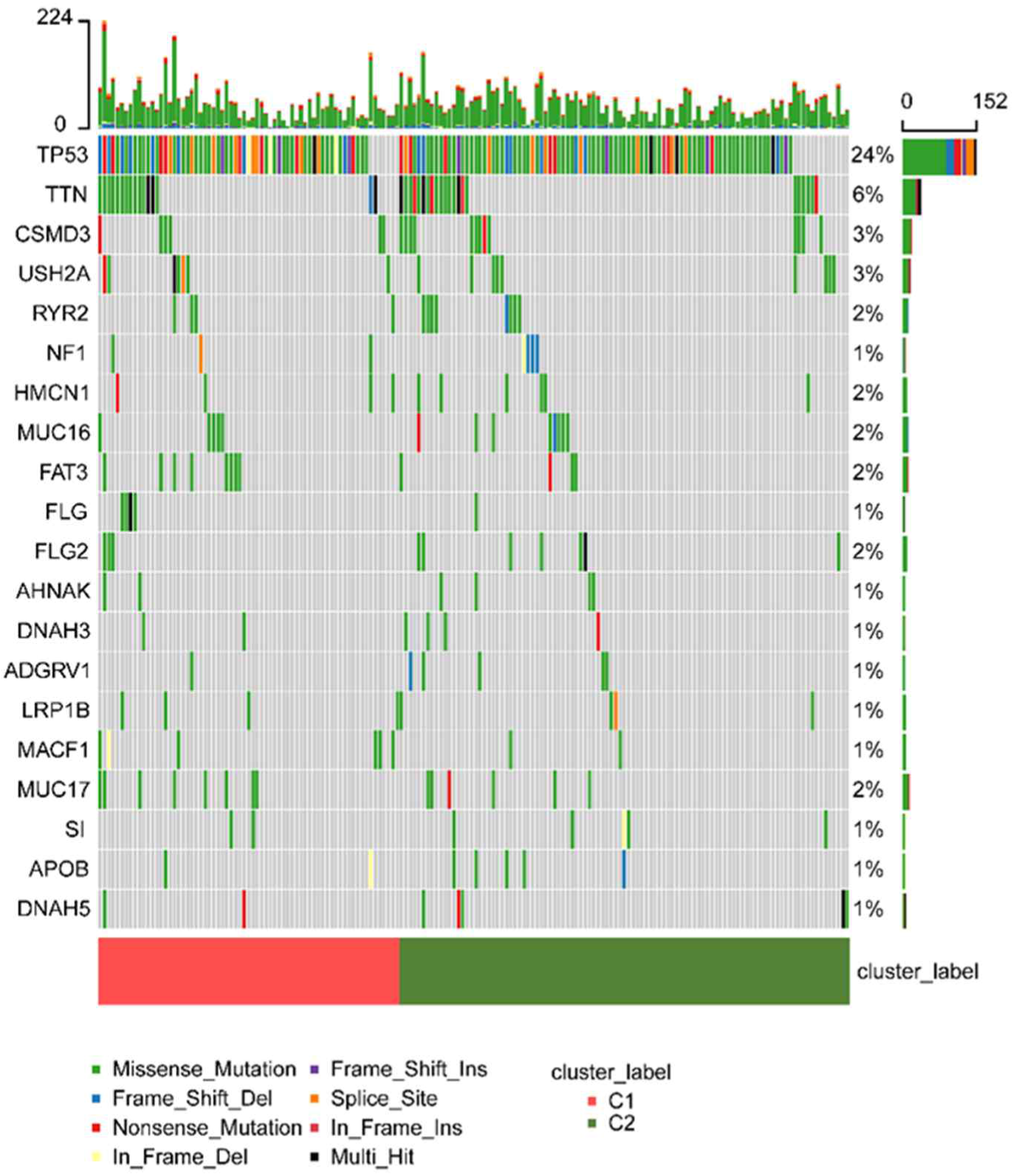
Mutational L-landscape of C1/C2-type samples in the TCGA cohort. Alterations in the top 20 most frequently mutated genes in TCGA-OV cohort. The mutation types and C1/C2 statuses are shown.

**Supplementary Figure 7.**
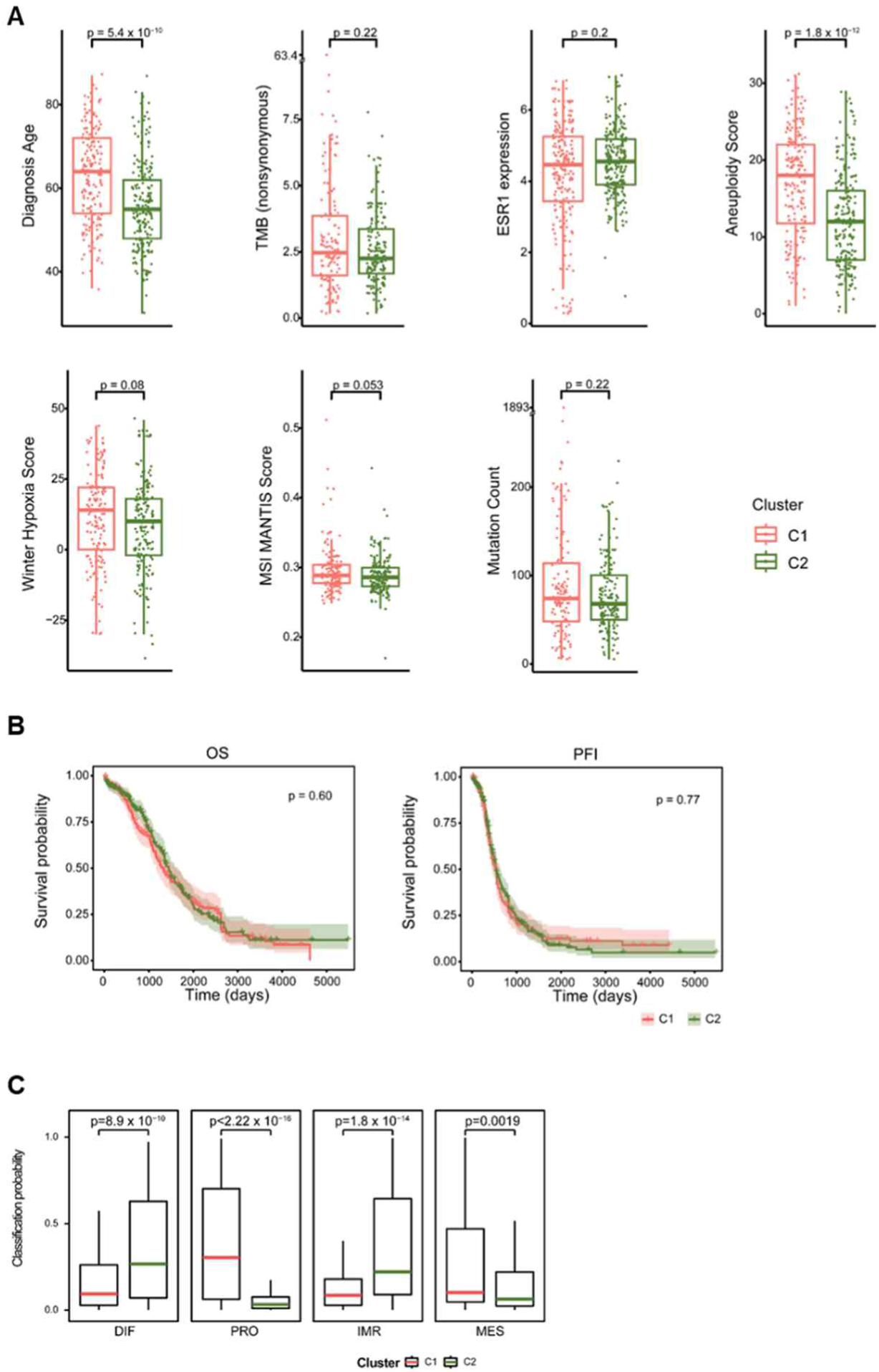
Clinical characteristics of C1/C2-type samples in TCGA cohort. A. Seven clinical characteristics of C1/C2-type samples obtained from cBioPortal: age at diagnosis, tumor mutation burden, ESR1 expression (log2[TPM+1]), aneuploidy score, winter hypoxia score, MSI score (MANTIS), and mutation count. B. Kaplan–Meier analysis of overall survival (OS) and progression-free interval (PFI) in C1- and C2-type samples. Statistical significance was calculated using log-rank tests. C. Classification probability of C1- and C2-type samples according to HGSOC consensus subtypes. Classification probability was calculated using the R package ConsensusOV. Box plots span the first to third quartiles and whiskers represent the 1.5 interquartile range. The central line indicates the median value. *P*-values were calculated using the Mann–Whitney U test.

**Supplementary Figure 8.**
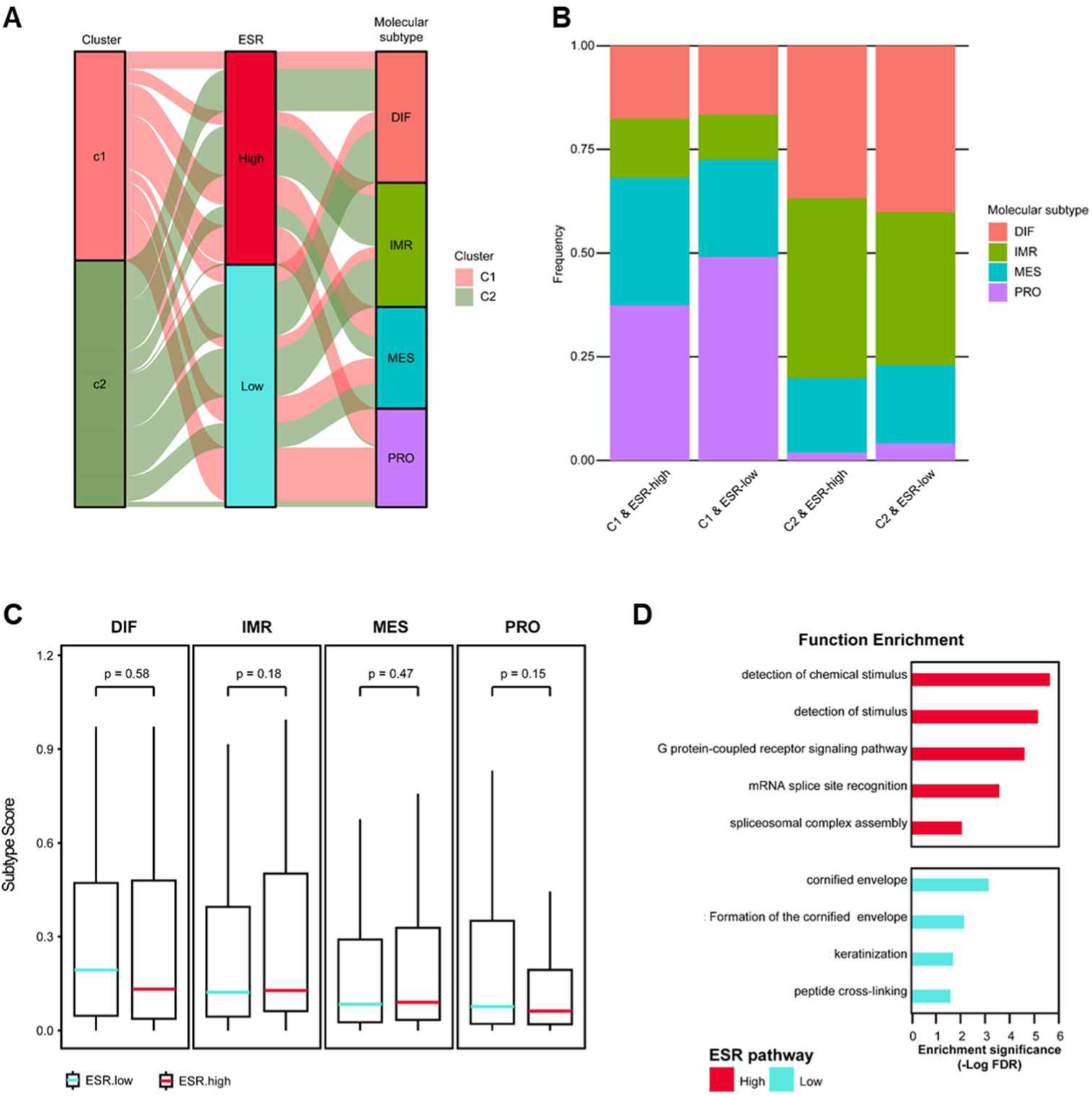
ESR signatures show no association with consensus molecular subtypes in the TCGA-OV cohort. a. Sankey plot illustrating the relationships between C1/2 subtypes, ESR signatures, and HGSOC consensus subtypes. b. Distribution of consensus subtypes across the four groups: C1 and ESR-high, C1 and ESR-low, C2 and ESR-high, and C2 and ESR-low. The threshold for ESR-high/low subtypes was defined as the mean value of the ESR-mediated signalling (R-HSA-8939211) enrichment score (ES). c. Classification probability of ESR-high (red line) and ESR-low (blue line) subtypes to HGSOC consensus subtypes. Box plots span the first to third quartiles and whiskers represent the 1.5 interquartile range. The central line indicates the median value. Statistical significance was calculated using the Mann–Whitney U test. d. Functional enrichment analysis results of DEGs between ESR-high (red) and ESR-low (sky blue) subtypes. Red bars represent enriched functions of the upregulated DEGs in ESR-high subtypes, whereas sky-blue bars represent enriched functions in ESR-low subtypes. Curated gene sets from the Gene Ontology, Reactome, and Transcription Factor databases were used.

**Supplementary Figure 9.**
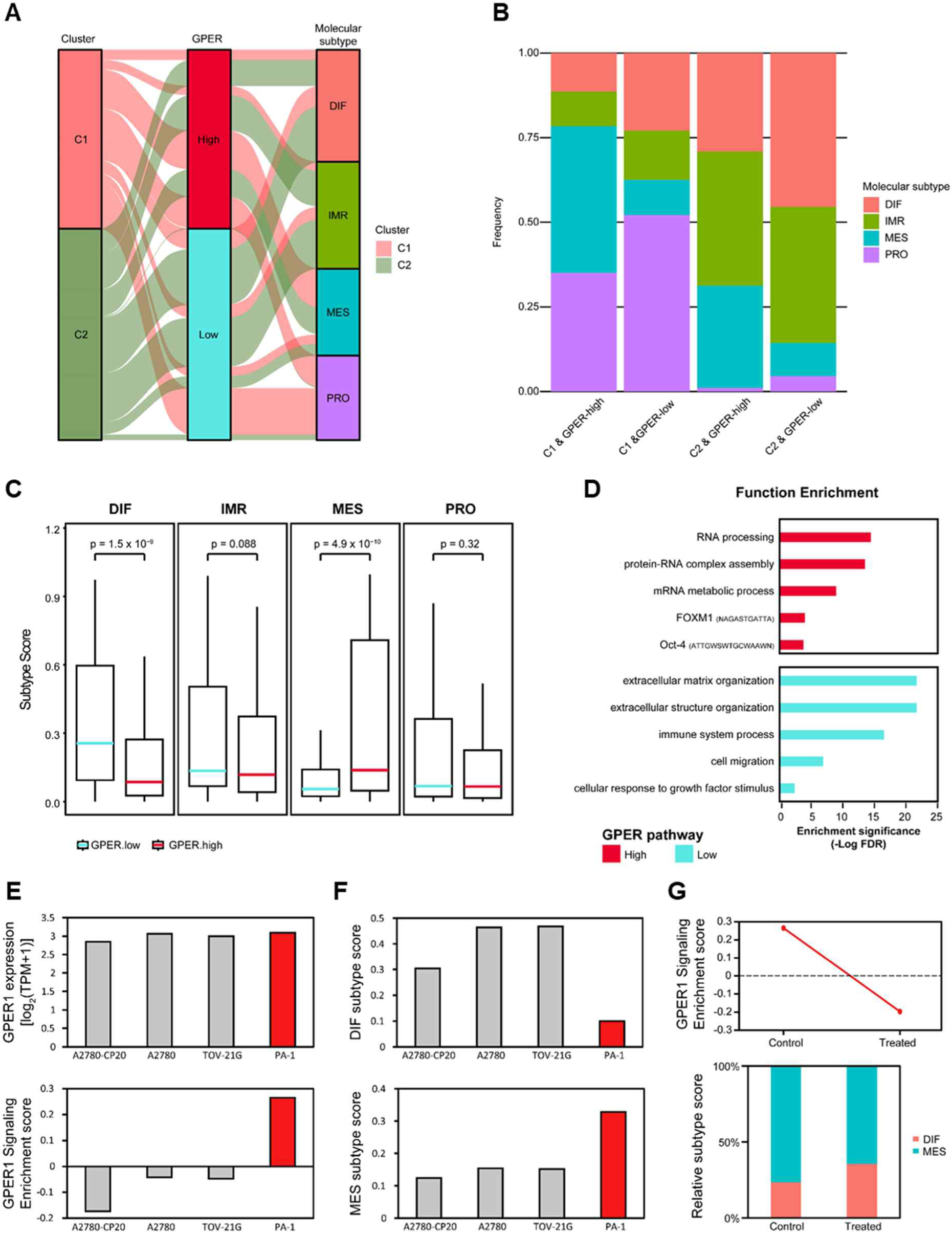
GPER signatures coordinate MES subtypes. A. Sankey plot showing the relationships between C1/2-type samples based on GPER1 signaling (R-HSA-9634597) and HGSOC consensus subtypes. B. Stacked bar plot showing the distribution of consensus subtypes across four groups: C1&GPER-high, C1&GPER-low, C2&GPER-high, and C2&GPER-low. The threshold for GPER high/low subtypes was defined as the mean GPER1 signaling (R-HSA-9634597) ES value. C. Classification probability of GPER-high (red) and GPER-low (sky blue) subtypes for HGSOC consensus subtypes. Box plots span from the first to third quartiles, and the whiskers represent the 1.5 interquartile range. The central line indicates the median value. Statistical significance was calculated using the Mann–Whitney U test. D. Functional enrichment analysis results of DEGs between GPER-high (red) and GPER-low (sky blue) subtypes. Curated gene sets from the Gene Ontology, Reactome, and TRANSFAC databases were used. E. Bar plots displaying GPER1 expression levels [log2(TPM+1)] (top) and ESs of the GPER1 signaling pathway (R-HSA-9634597, bottom) in the indicated ovarian cancer cell lines. F. Bar plots show subtype scores for DIF (top) and MES (bottom) across the indicated ovarian cancer cell lines. Subtype scores represent the classification probabilities calculated using the R package ConsensusOV. G. ES of the GPER1 signaling pathway in PA-1 cells treated or not (control) with G15 (GPER1 inhibitor) (50 μM, 12 h, top). Stacked bar plot showing relative subtype scores between the DIF and MES subtypes in the control and G15-treated cells (bottom).

**Supplementary Figure 10.**
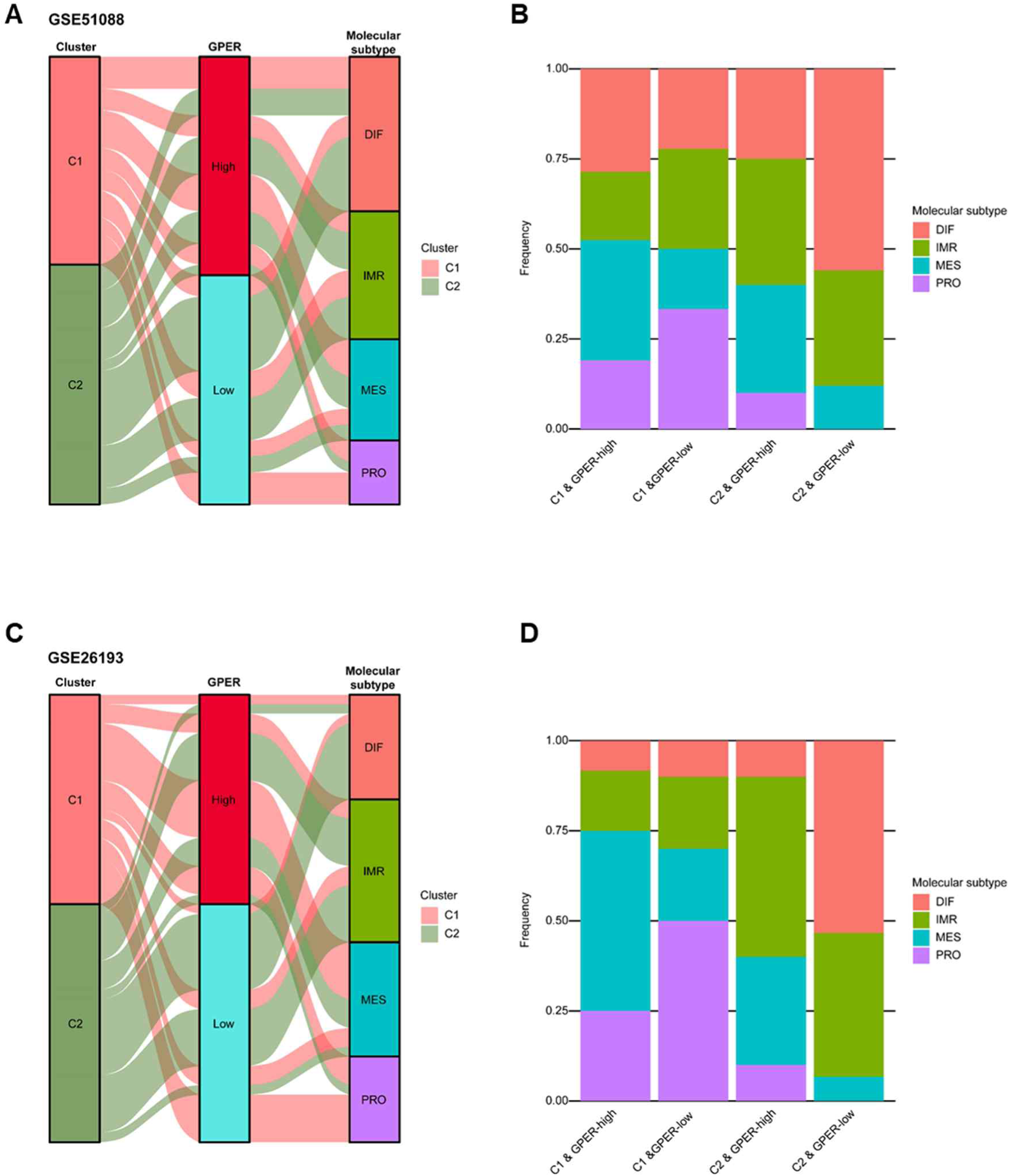
Association of GPER signatures with mesenchymal subtype-related genes in two ovarian cancer cohorts (GSE51088 and GSE26193). A, **C** Sankey plots illustrating the relationships among C1/2 type samples, GPER signatures, and HGSOC consensus subtypes in the GSE51088 (A) and GSE26193 (C) cohorts. **B, D** Distribution of consensus subtypes across the four groups in GSE51088 (B) and GSE26193 (D) cohorts: C1 and GPER-high, C1 and GPER-low, C2 and GPER-high, and C2 and GPER-low. The threshold for the GPER-high/low subtypes was defined as the mean value of GPER1 signalling (R-HSA-9634597) ES.

**Supplementary Figure 11.**
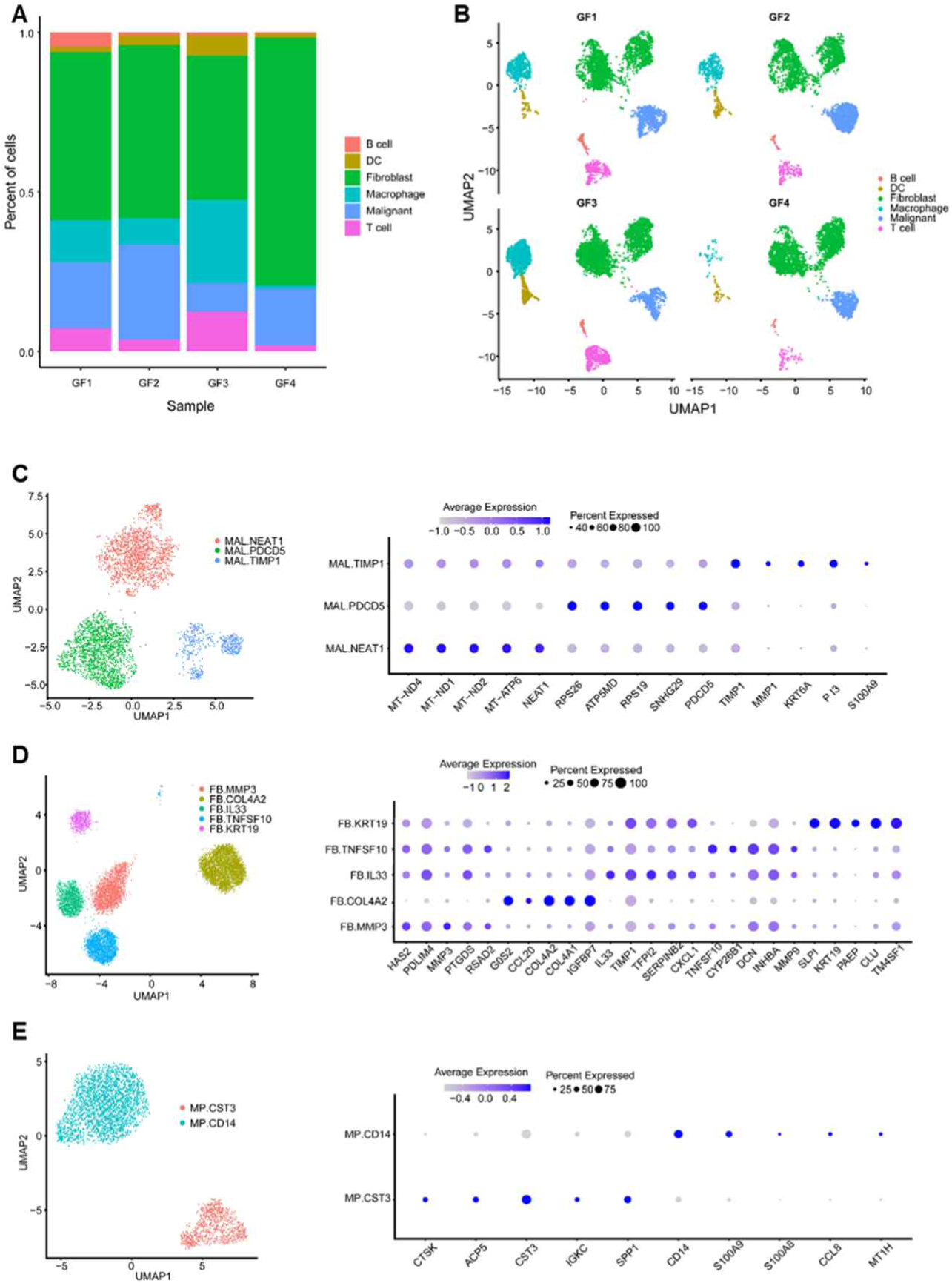
General single-cell characteristics of four ovarian cancer samples. A. Distribution of cell populations across four ovarian cancer samples in the presence or absence of estradiol based on single-cell RNA sequencing. B. Uniform Manifold Approximation and Projection plot showing cell populations across four ovarian cancer samples. C–E. Subtypes of malignant cells (C), fibroblasts (D), and macrophages (E) were identified using PRIMUS analysis. The subtyping data are summarized in UMAP plots and color-coded according to the cell subtypes (left). The top five markers for each subtype are shown as dotted plots (right panels). Color bars indicate the average expression of genes in the subtypes, and the dot size represents the percentage of cells expressing these genes. Markers were selected based on the fold change between cells.

**Supplementary Figure 12.**
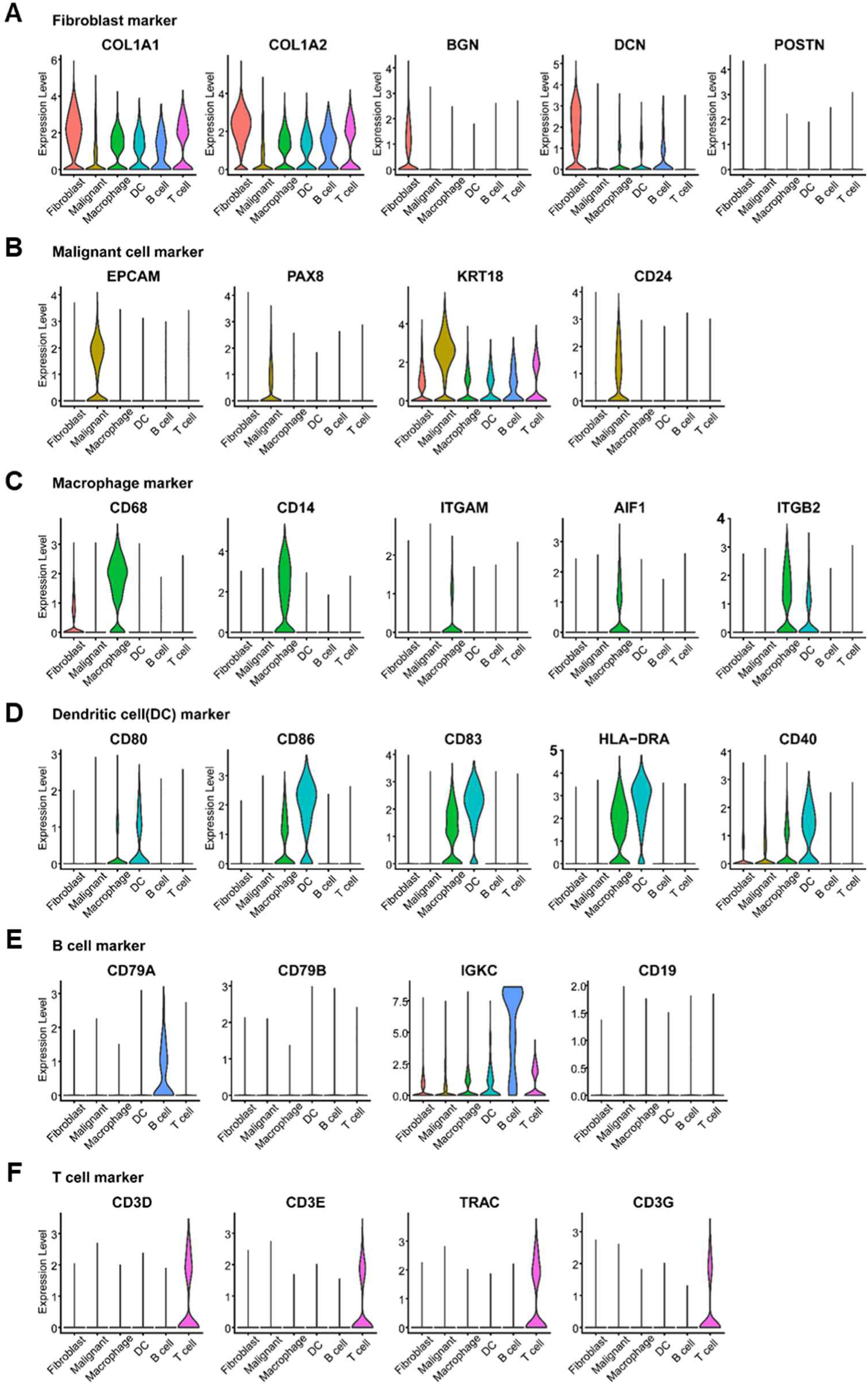
Expression pattern of major cell markers that define the distinct cell fractions. a–f. Violin plots showing representative marker expression of the indicated cell types: fibroblasts (A), malignant cells (B), macrophages (C), dendritic cells (D), B cells (E), and T cells (F).

**Supplementary Figure 13.**
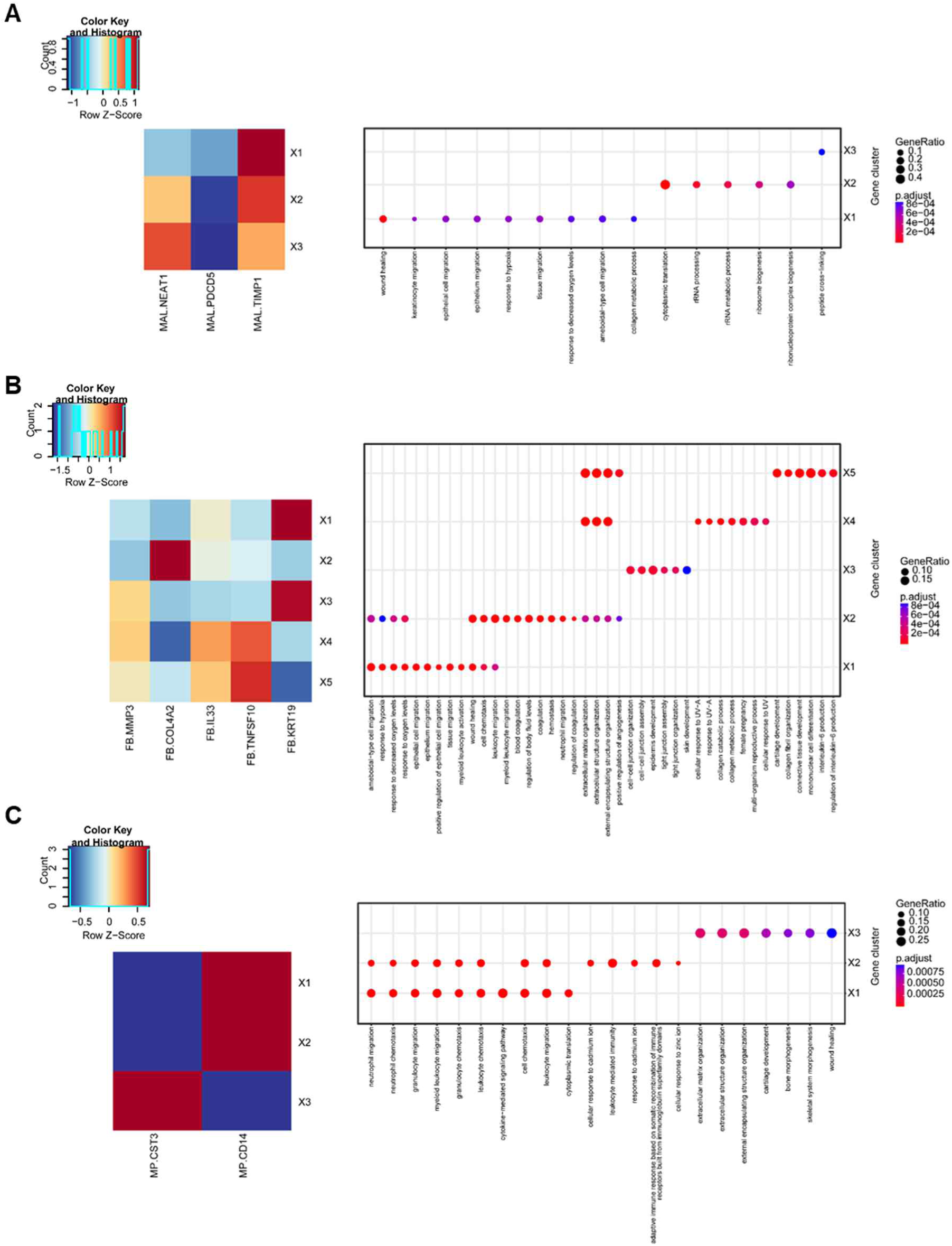
Representative gene sets of major cell clusters in single-cell analysis. A–C. Heatmaps (left) and dot blots (right) show the normalized ESs of the indicated cell subtypes for malignant cells (A), fibroblasts (B), and macrophages (C). Dot color indicates the adjusted *P*-value of enrichment, and dot size represents the overlap between representative gene sets and curated gene sets from Gene Ontology.

**Supplementary Figure 14.**
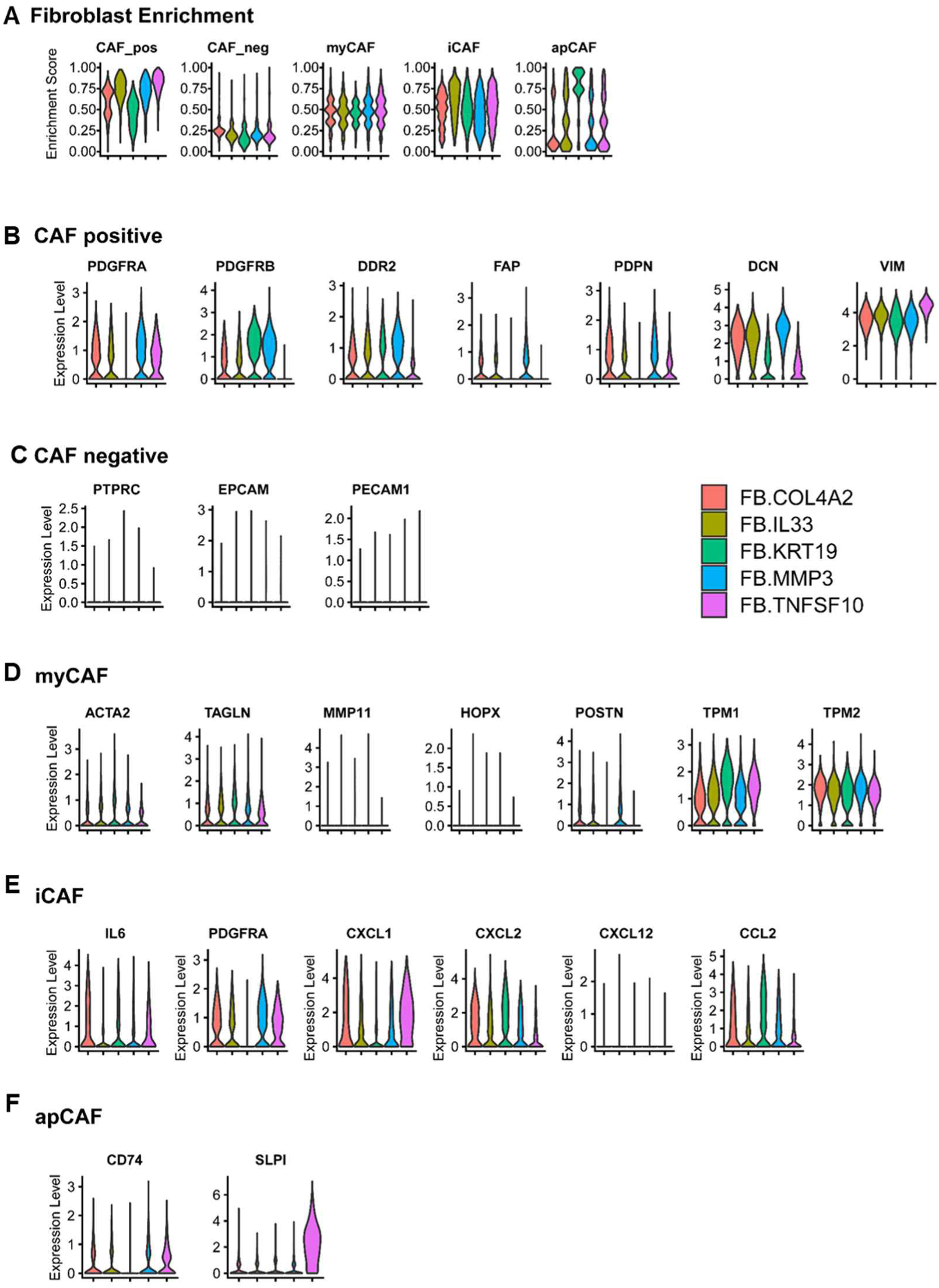
Analysis of CAF subtype signatures in single-cell fibroblast fractions. A. ESs of five fibroblast subtypes for CAF markers and subtype markers. B, C. Gene expression levels of CAF-positive and -negative markers. D–F. Gene expression levels in three CAF subtypes: myofibroblastic CAFs (myCAF), inflammatory CAFs (iCAF), and antigen-presenting CAFs (apCAF).

**Supplementary Figure 15.**
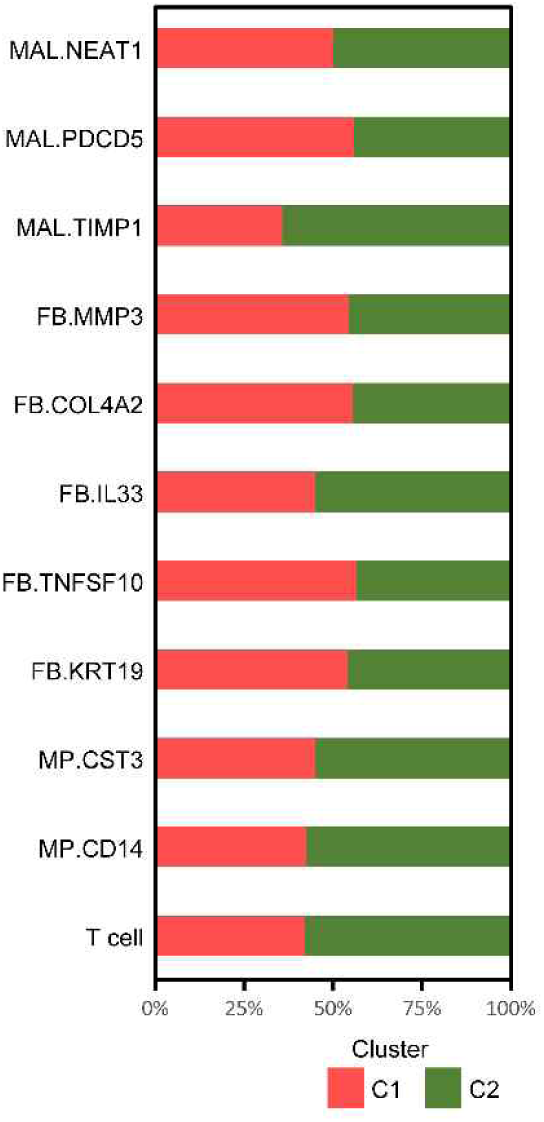
Relative fractions of cell subtypes in TCGA-OV samples. The indicated cell fractions were determined by CIBERSORTx cell deconvolution of C1-(red) and C2-(green)-type samples.

**Supplementary Figure 16.**
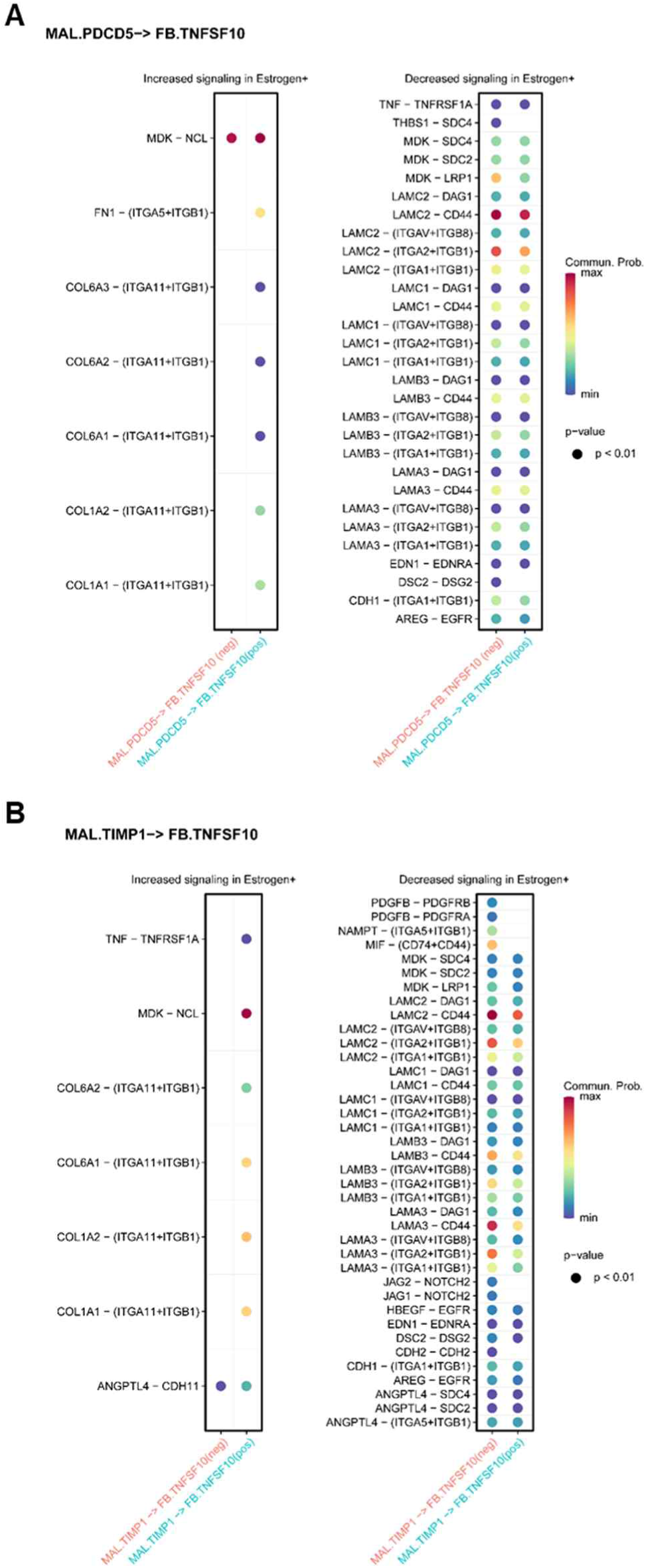
Ligand–receptor interaction between malignant cells and fibroblasts. Differential communication probability of ligand–receptor pairs that increased (left) or decreased (right) in estradiol-treated (sky blue) versus non-treated (pink) conditions between MAL.PDCD5 and TNFSF10 (A) and between MAL.TIMP1 and FB.TNFSF10 (B).

**Supplementary Figure 17.**
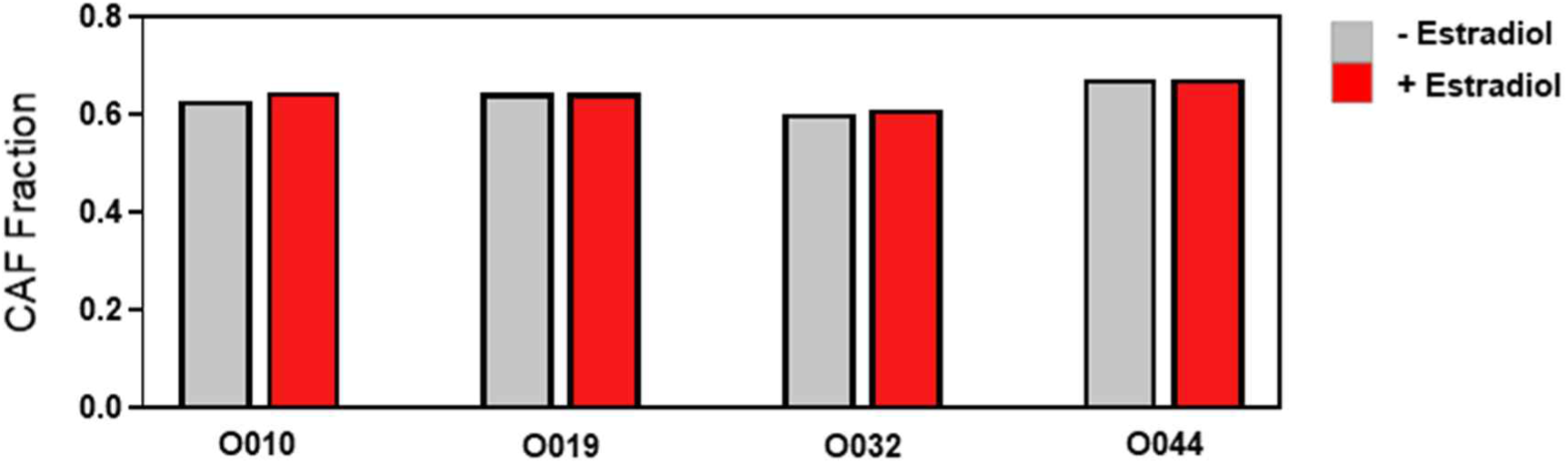
Analysis of CAF fractions in established CAF lines. Bar plots show CAF fractions calculated using CIBERSORTx in established CAFs under estradiol-treated and estradiol-untreated conditions.

**Supplementary Figure 18.**
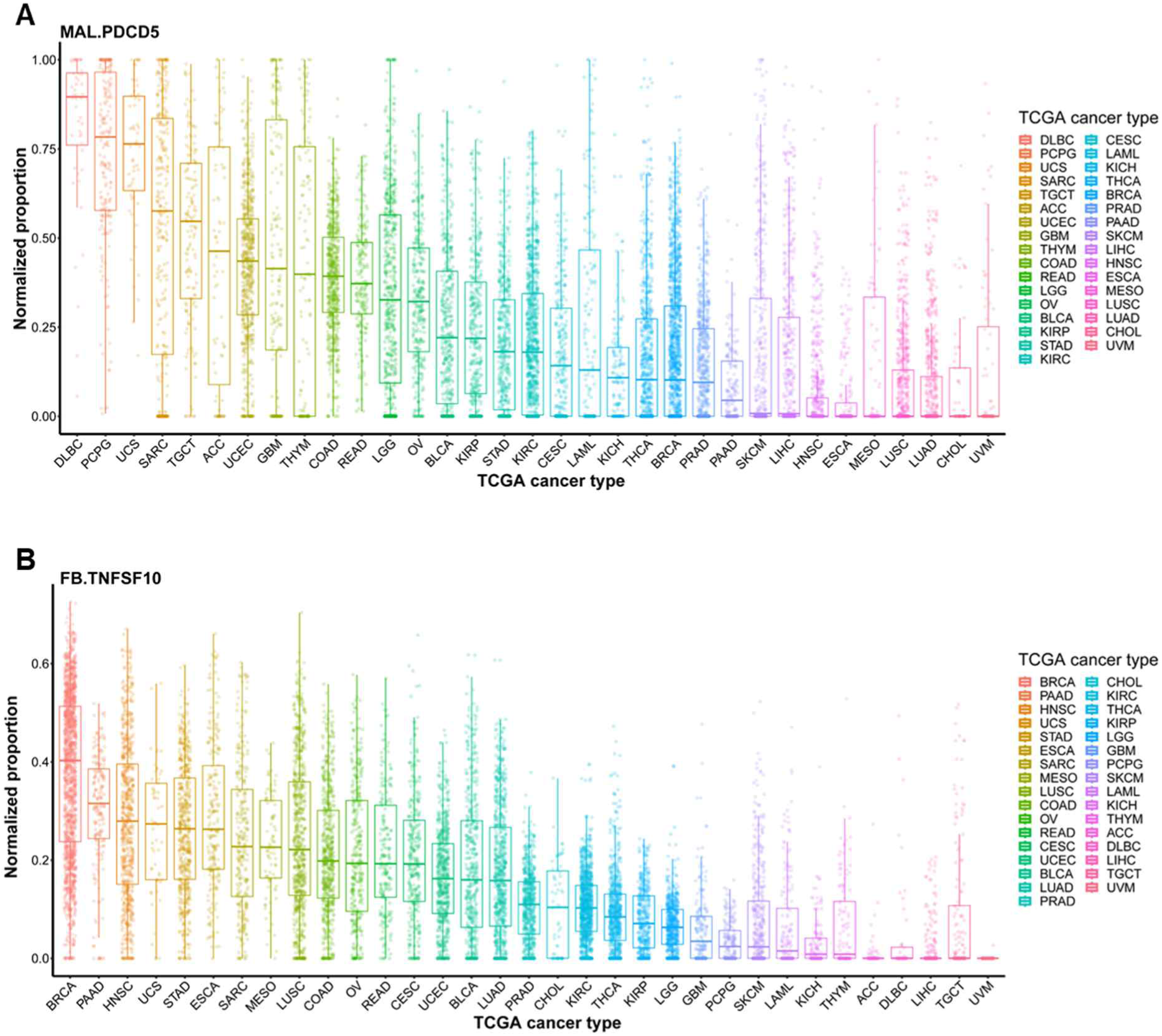
The cell deconvolution results of FB.TNFSF10 and MAL.PDCD5 cells across 33 TCGA cancer types. A, B. The normalized proportions of the two subtypes, FB.TNFSF10 (A) and MAL.PDCD5 (B), were measured across 33 TCGA cancer types using CIBERSORTx.

**Supplementary Figure 19.**
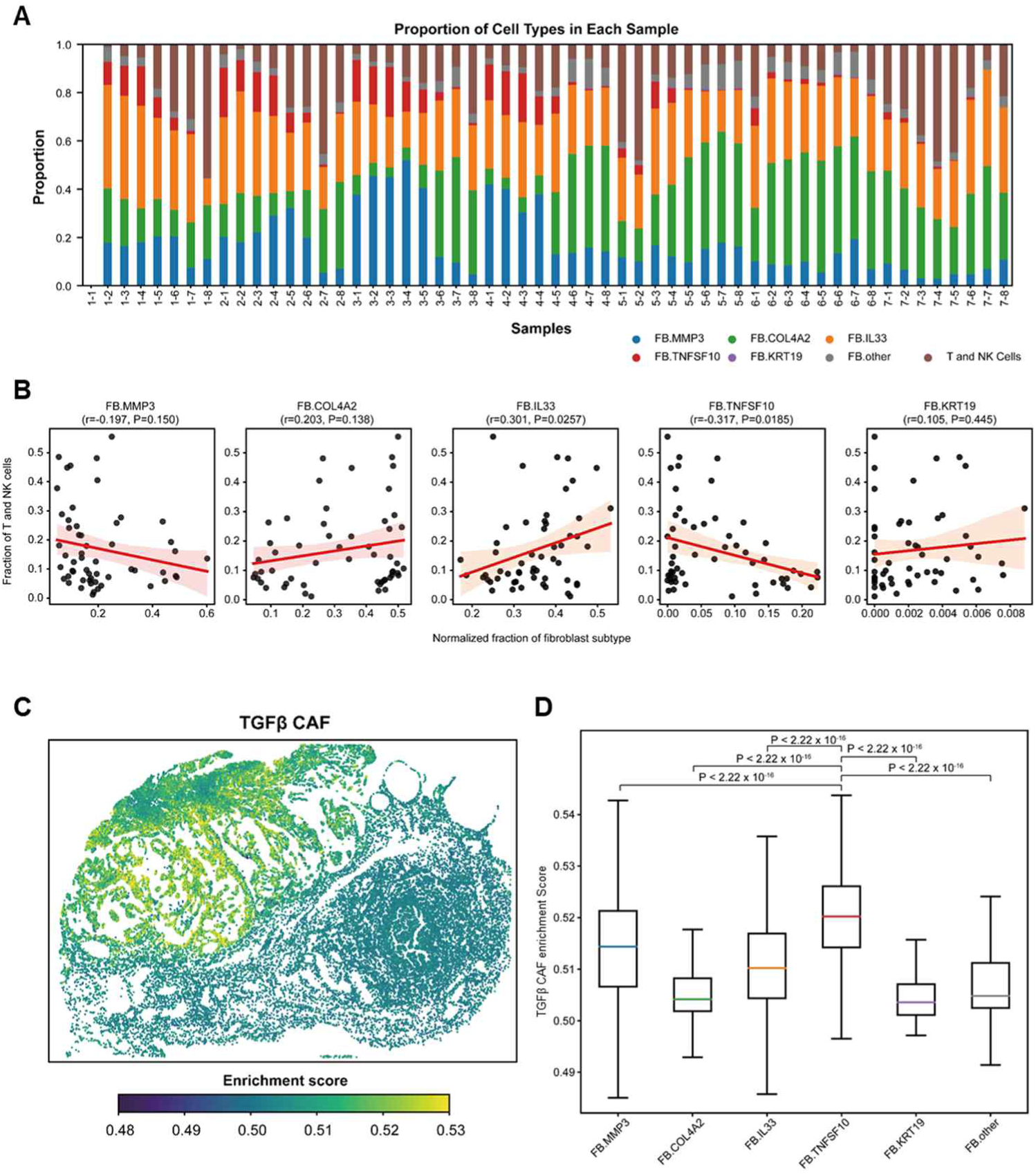
Spatial associations between FB fractions and immune infiltration. A. The Xenium dataset was divided into a 7×8 grid (56 tiles, labeled S1-1 through S7-8), and cell type proportions were quantified for each tile. Stacked bar plots show the relative abundance of different FB subtypes and T/NK cells. B. Correlation analysis between normalized FB subtype proportions and T/NK cell fractions across the tiles. Pearson’s correlation coefficients (PCC) were calculated for each fibroblast subtype. Black dots represent individual tiles (n = 56) and the red line indicates the linear regression fit with a 95% confidence interval (shaded area). **C**, TGF-β CAF enrichment scores in a Xenium data (left). Box plots show the quartiles and 1.5 IQR whiskers. The median is marked by the central line. Significance was assessed using the Mann-Whitney U test (right).

## Notes

All authors declare no conflict of interest.

### Competing Interest Statement

The authors have declared no competing interest.

